# 1700029I15Rik orchestrates the biosynthesis of acrosomal membrane proteins required for sperm–egg fusion

**DOI:** 10.1101/2022.04.15.488448

**Authors:** Yonggang Lu, Kentaro Shimada, Jingjing Zhang, Yo Ogawa, Shaogeng Tang, Taichi Noda, Hiroki Shibuya, Masahito Ikawa

## Abstract

Sperm acrosomal membrane proteins, such as IZUMO1 and SPACA6, play an essential role in mammalian sperm–egg fusion. How their biosynthesis is regulated during spermiogenesis has largely remained unknown. Here, we show that the *1700029I15Rik* knockout male mice are severely subfertile and their spermatozoa do not fuse with eggs. *1700029I15Rik* encodes a type-II transmembrane protein that is expressed in early spermatids but not in mature spermatozoa. 1700029I15Rik is associated with proteins involved in N-glycosylation, disulfide isomerisation, and ER– Golgi trafficking, suggesting its involvement in nascent protein processing. *1700029I15Rik* knockout testis has a normal level of sperm plasma membrane proteins, but decreased expression of multiple acrosomal membrane proteins. The knockout sperm exhibit elevated ubiquitinated proteins and upregulated ER-associated degradation; strikingly, SPACA6 becomes undetectable. Our results support for a specific, 1700029I15Rik-mediated pathway in spermiogenesis for the assembly of acrosomal membrane proteins.

**Significance Statement:** In sexually reproducing species, life begins with the fusion between a sperm and an egg. Multiple sperm acrosomal membrane proteins have been reported indispensable for sperm–egg fusion in mammals, yet the mechanism underlying their biosynthesis remains unknown. The present study demonstrates the existence of a 1700029I15Rik-mediated pathway specifically coordinating the processing and assembly of acrosomal membrane proteins. It represents an intriguing paradigm where the biosynthesis of proteins destined for various subcellular compartments might be orchestrated in a spatiotemporal manner. Given 1700029I15Rik is highly conserved in human, our findings provide potential insights into the aetiology of idiopathic male infertility and the development of a novel contraceptive approach involving molecular interventions in the maturation of gamete fusion-required acrosomal proteins.

## Introduction

Mammalian fertilisation is a complicated series of events culminating in the union of a haploid spermatozoon and a haploid egg to produce a diploid zygote. This process is a fundamental but obligatory prerequisite for successful propagation of parental genomes to the next generation. During the journey through the female reproductive tract, spermatozoa undergo the acrosome reaction to expose the acrosomal contents, including hydrolytic enzymes and acrosomal membrane proteins, to gain competence in penetrating the egg zona pellucida (ZP) and fusing with the oolemma (1).

The plasma membrane fusion between gametes is a fascinating eukaryotic cell–cell fusion event involving the merger of two morphologically distinct cells from individuals of opposite sexes. Notwithstanding the decades-long research effort, detailed cellular and molecular mechanisms underlying this unique fusion process remain shrouded in mystery. To date, seven sperm proteins, IZUMO1 (2), SPACA6 (3–5), FIMP (6), SOF1 (4), TMEM95 (4, 7), and DCST1 and DCST2 (8, 9), have been demonstrated essential for sperm–egg fusion. The knockout mouse sperm lacking any of these molecules fail to fuse with the oolemma, despite their normal morphology, motility, and the ability to elicit the acrosome reaction. Among these proteins, IZUMO1, SPACA6, TMEM95, DCST1, and DCST2 are transmembrane glycoproteins that initially localise to the acrosomal membrane and translocate to the sperm surface during the acrosome reaction (2, 4, 9, 10).

In the early spermatids, the acrosomal precursor proteins are synthesised and processed in the endoplasmic reticulum (ER), trafficked to the Golgi apparatus, and secreted as proacrosomal vesicles that subsequently coalesce to form the acrosome (11). In this study, by harnessing genomics, proteomics, and cell biology techniques, we unravel that 1700029I15Rik, an ER-expressing type-II transmembrane protein, facilitates the biosynthesis of acrosomal membrane proteins involved in sperm–egg fusion.

## Results

### 1700029I15Rik is a testis-specific type-II transmembrane protein expressed during early spermiogenesis

Mouse *1700029I15Rik* encompasses three coding exons and is localised to the forward strand of chromosome 2 (SI Appendix, Fig. S1a). This gene localises to chromosome 11 in human, thus named as *C11orf94*. RT-PCR revealed the expression of *1700029I15Rik* exhibited a bias towards testis, and initiated at postnatal day 21 (Fig. 1a, b), corresponding to the early spermiogenesis in the first wave of mouse spermatogenesis. Consistently, a previously published single-cell RNA-sequencing (scRNA-seq) analysis (12) indicated *1700029I15Rik* was expressed in the mid-round spermatids (SI Appendix, Fig. S1b). 1700029I15Rik is highly conserved among mammalian species (Fig. 1c<u>, d</u> and SI Appendix, Fig. S1c). In certain species, such as black flying fox and green sea turtle, 1700029I15Rik is absent but PEX16, a peroxisomal membrane protein, carries a C-terminal region that shows high homology with 1700029I15Rik (SI Appendix, Fig. S1d – f). Given *Pex16* is localised immediately upstream of *1700029I15Rik* in the genome (SI Appendix, Fig. S1a), we speculate that these species harbour a *Pex16*-*1700029I15Rik* chimeric fusion transcript.

**Fig. 1.**
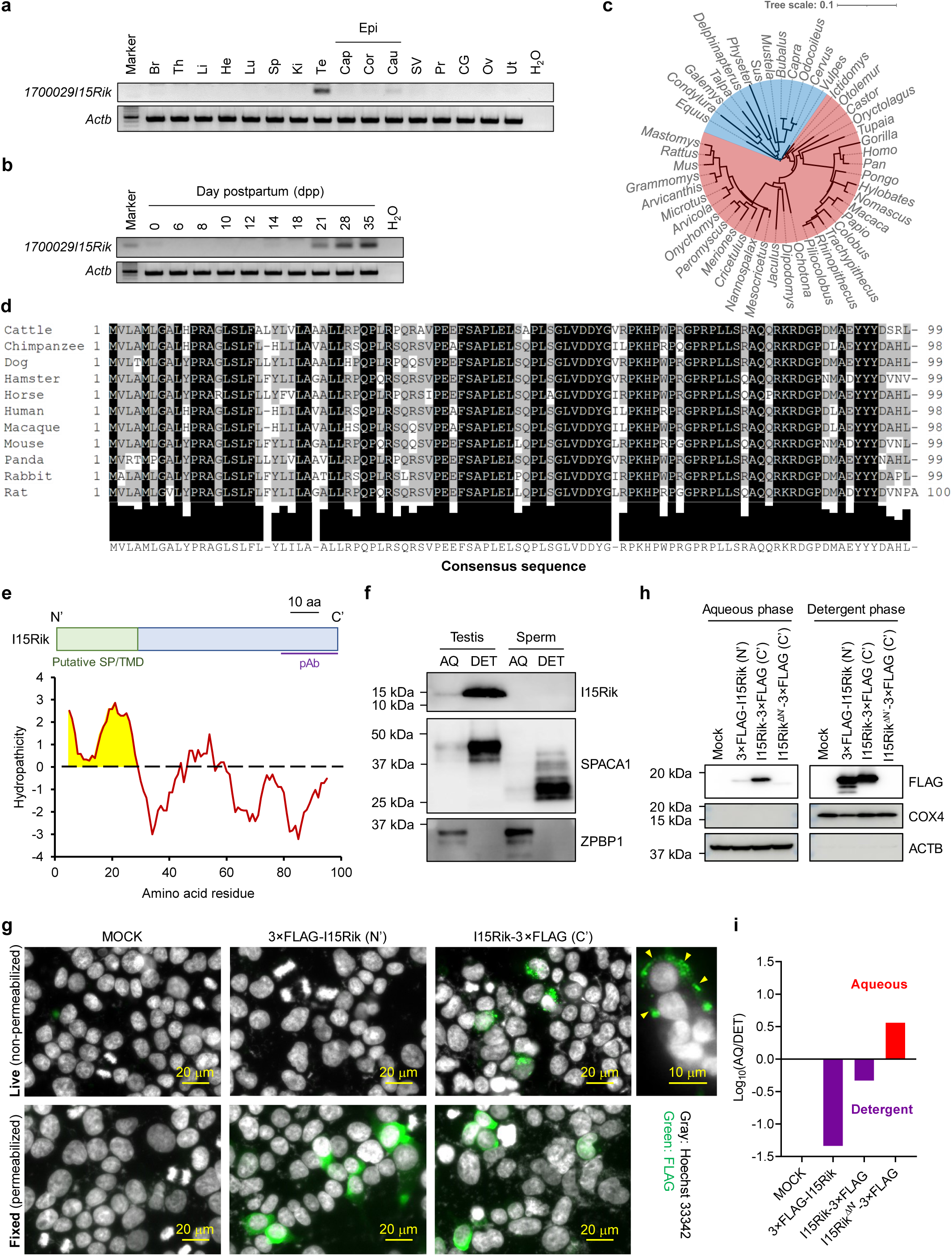
*1700029I15Rik* is a testis-enriched type-II transmembrane protein conserved in mammals and expressed during spermiogenesis. **a**, Analyses of mRNA expression of *1700029I15Rik* in various mouse tissues by RT-PCR. Br, brain; Th, thymus; Li, liver; He, heart; Lu, lung; Sp, spleen; Ki, kidney; Te, testis; Cap, caput epididymis; Cor, corpus epididymis; Cau, cauda epididymis; Epi, epididymis; SV, seminal vesicle; Pr, prostate; CG, coagulating gland; Ov, ovary; Ut, uterus. The expression of *β-actin* (*Actb*) was analysed as a loading control. **b**, Analyses of mRNA expression of *1700029I15Rik* in mouse testes during postnatal development. **c**, Phylogenetic tree indicating the evolutionary conservation of 1700029I15Rik in various mammals. The tree was visualised by the iTOL software (27). Red and blue highlighted species belong to Euarchontoglires and Laurasiatheria, respectively. **d**, Multiple sequence alignment of 1700029I15Rik proteins in 11 mammalian species. The lower panel indicates the consensus sequence and the extent of sequence conservation among the mammalian species. **e**, A predicted structure and hydropathy analysis of 1700029I15Rik. The N-terminus of 1700029I15Rik is predicted as a signal peptide (SP) or a transmembrane domain (TMD). A polyclonal antibody (pAb) was generated against the C-terminal region. Hydropathicity of each amino acid was determined by the Kyte-Doolittle scale (28). The hydrophobic region highlighted in yellow corresponds to the putative SP/TMD. **f**, Triton X-114 fractionation of testis and sperm proteins. SPACA1 and ZPBP1 were analysed as positive controls for proteins in the aqueous (AQ) and detergent (DET) phases, respectively. **g**, Topological analysis of 1700029I15Rik in vitro by live cell immunostaining. Yellow arrowheads indicate fluorescence signals at the cell surface. **h – i**, Analysis of the subcellular localisation of 1700029I15Rik before and after deletion of the N-terminal hydrophobic region. The band intensities were analysed by ImageJ and relative expression was calculated by Log_10_(AQ/DET).

1700029I15Rik contains a hydrophobic N-terminus [amino acids (aa) 1 – 29] predicted as either a transmembrane helix or a signal peptide (Fig. 1e and SI Appendix, Table S1). For further analyses, we produced a polyclonal antibody against its evolutionarily conserved C-terminal region (Fig. 1e and SI Appendix, Fig. S1g). Triton X-114 subcellular fractionation unveiled that 1700029I15Rik is a membrane-bound protein exclusively detected in the detergent phase of testis lysate (Fig. 1f). To analyse the protein topology, we constructed plasmids encoding N-terminal or C-terminal 3×FLAG-tagged 1700029I15Rik (SI Appendix, Fig. S1h, i). Live cell immunostaining indicated that only the HEK293T cells expressing the C-terminal tagged 1700029I15Rik exhibited fluorescence signals on the cell surface (Fig. 1g), suggesting the C-terminus is exposed extracellularly. Notably, Phyre2 predicts another hydrophobic region (aa 46 – 61) as a transmembrane domain (SI Appendix, Table S1 and Fig. 1e). To examine this possibility, we further generated a plasmid carrying 1700029I15Rik lacking the N-terminal hydrophobic region (aa 1 – 29; SI Appendix, Fig. S1h). The truncated 1700029I15Rik was detected only in the aqueous phase, indicating the second hydrophobic region does not span the membrane (Fig. 1h, i). Taken together, these experiments demonstrate 1700029I15Rik is a type-II transmembrane protein that anchors to the membrane via its N-terminus.

### *1700029I15Rik* knockout male mice are infertile due to defective sperm–egg fusion

To examine the function of *1700029I15Rik* in vivo, we generated a knockout mouse line on the background of B6D2F1 by CRISPR/Cas9 (Fig. 2a and SI Appendix, SI Materials and Methods). A 619 bp-deletion was introduced to the coding region of *1700029I15Rik*, resulting in the translation of an incorrect amino acid sequence (Fig. 2a, b and SI Appendix, Fig. S2a). We confirmed the absence of 1700029I15Rik in the knockout testis by western blot. 1700029I15Rik was not detected in the wildtype sperm lysate (Fig. 1f, 2c), implying this protein functions during spermatogenesis.

**Fig. 2.**
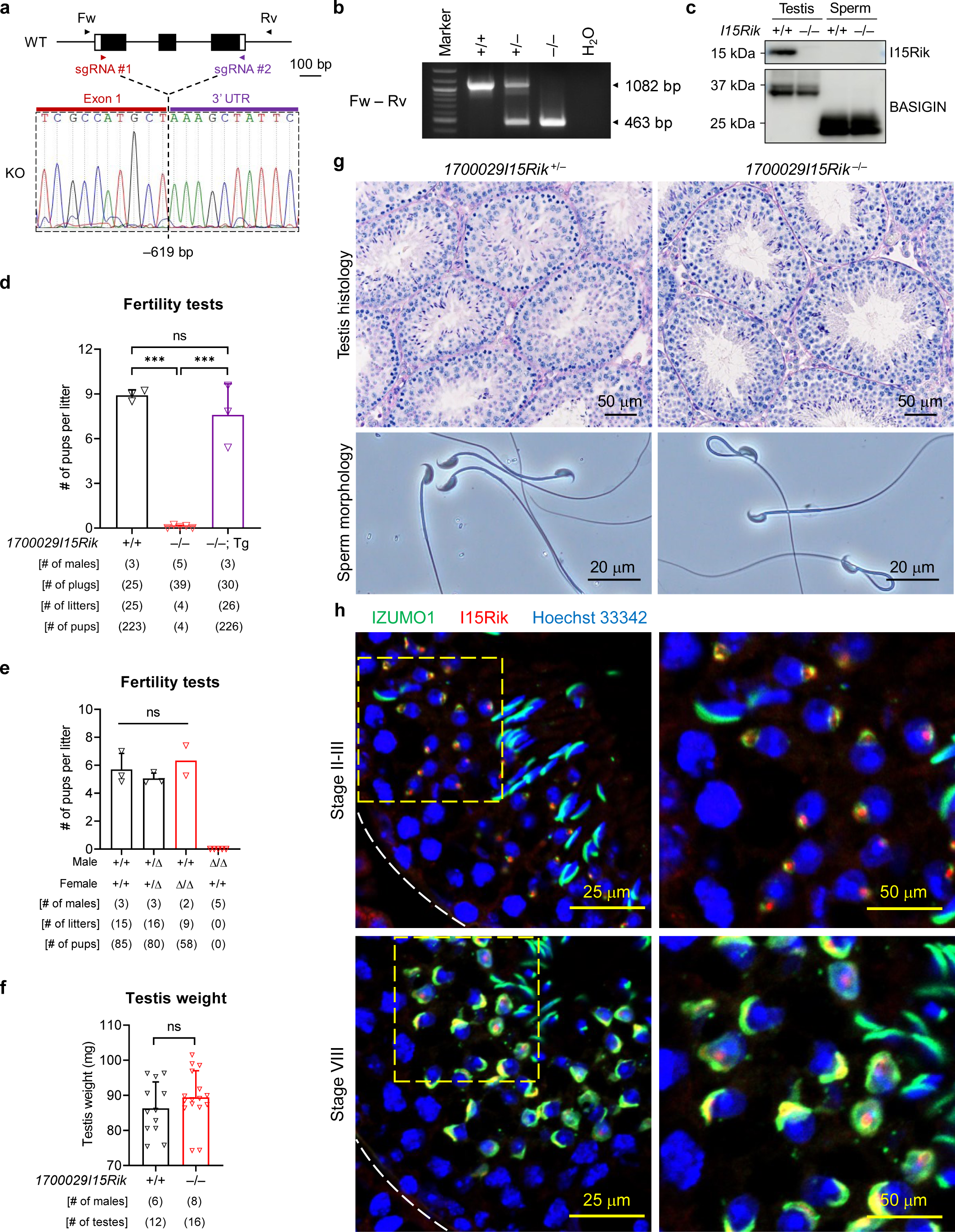
*1700029I15Rik* knockout males are infertile despite normal spermatogenesis and sperm morphology. **a**, Schematic representation of the knockout strategy of *1700029I15Rik* in mice by CRISPR/Cas9. Single guide RNAs (sgRNAs) #1 and #2 were designed to target the first coding exon and the 3’ untranslated region (UTR), respectively. **b**, Genomic PCR for distinguishing the wildtype, and heterozygous and homozygous knockout offspring using the forward (Fw) and reverse (Rv) primers. **c**, Immunodetection of 1700029I15Rik in the wildtype and knockout testis and sperm. BASIGIN was analysed in parallel as a loading control. **d**, Fertility tests of *1700029I15Rik*^+/+^, *1700029I15Rik*^−/−^, and *1700029I15Rik*^−/−^; Tg males. ns, not significant. **e**, Fertility tests of *1700029I15Rik*^Δ/Δ^ mice. **f**, Comparison of the testis weights in *1700029I15Rik*^+/+^ and *1700029I15Rik*^−/−^ males. **g**, Analyses of testis histology and sperm morphology in *1700029I15Rik*^+/+^ and *1700029I15Rik*^−/−^ males. **h**, Co-immunostaining of IZUMO1 and 1700029I15Rik in the wildtype testis cryosections.

In parallel, we produced *1700029I15Rik* mutant mice lacking 215 bp at the first coding exon on a pure C57BL/6J background (SI Appendix, Fig. S2b<u>, c</u> and SI Materials and Methods). Both the knockout (*1700029I15Rik*^−/−^) and mutant (*1700029I15Rik*^Δ/Δ^) mice exhibited normal development, appearance, and behaviours. *1700029I15Rik*^−/−^ female mice exhibited normal fecundity, whereas *1700029I15Rik*^−/−^ males were severely subfertile despite successful coituses (SI Appendix, Fig. S2d – f). A C-terminal PA-tagged *1700029I15Rik* transgene (Tg) restored the fecundity of knockout males (Fig. 2d and SI Appendix, Fig. S2g – i), demonstrating the male infertility was not derived from off-target mutations. *1700029I15Rik*^Δ/Δ^ males were sterile (Fig. 2e), indicating the phenotype severity correlates with the genetic backgrounds. The testis weight, spermatogenesis, and sperm morphology were normal in *1700029I15Rik*^−/−^ males (Fig. 2f, g). Likewise, *1700029I15Rik*^Δ/Δ^ males showed no abnormalities in the testis and epididymis histology, as well as the first meiotic division of spermatocytes (SI Appendix, Fig. S2j– l). To avoid unnecessary duplication of efforts, all subsequent studies were performed on the *1700029I15Rik* knockout mouse line.

1700029I15Rik initially appeared at the centre of the proacrosomal vacuole in step 2 – 3 spermatids, distinct to IZUMO1 that localised to the proacrosomal membrane (Fig. 2h and SI Appendix, Fig. S3a). In step 8 spermatids, 1700029I15Rik was partially co-localised with IZUMO1 in the proacrosome. Nevertheless, 1700029I15Rik, but not IZUMO1, was detected in the acrosomal granule (Fig. 2h). From step 9 spermatids, 1700029I15Rik signals were disappeared from the acrosome and translocated to the manchette (SI Appendix, Fig. S3a). The manchette staining was non-specific, because the same staining pattern was observed in the knockout spermatids (SI Appendix, Fig. S3b). These observations unravel a unique subcellular localisation of 1700029I15Rik in the early spermatids.

*1700029I15Rik* knockout spermatozoa showed significantly reduced ability to fertilise the cumulus-intact, cumulus-free, and ZP-free eggs in vitro (Fig. 3a and SI Appendix, Fig. S4a, b). In vivo, the knockout sperm failed to fertilise the eggs and were accumulated in the perivitelline space, indicating the sperm underwent the acrosome reaction and penetrated the ZP but failed to fuse with the oolemma (Fig. 3b, c and SI Appendix, Fig. S4c). Through sperm–oolemma fusion and binding assays, we confirmed that *1700029I15Rik* knockout sperm could bind to but not fuse with the oolemma (Fig. 3d – g). Despite the impaired fertilising ability, the knockout sperm exhibited normal motility, velocity, and ability to bind to the ZP (SI Appendix, Fig. S4d – h). Furthermore, HEK293T cells transiently expressing 1700029I15Rik did not bind to or fuse with the ZP-free mouse eggs (SI Appendix, Fig. S4i, j). These findings suggest 1700029I15Rik is essential for, but does not directly participate in gamete fusion.

**Fig. 3.**
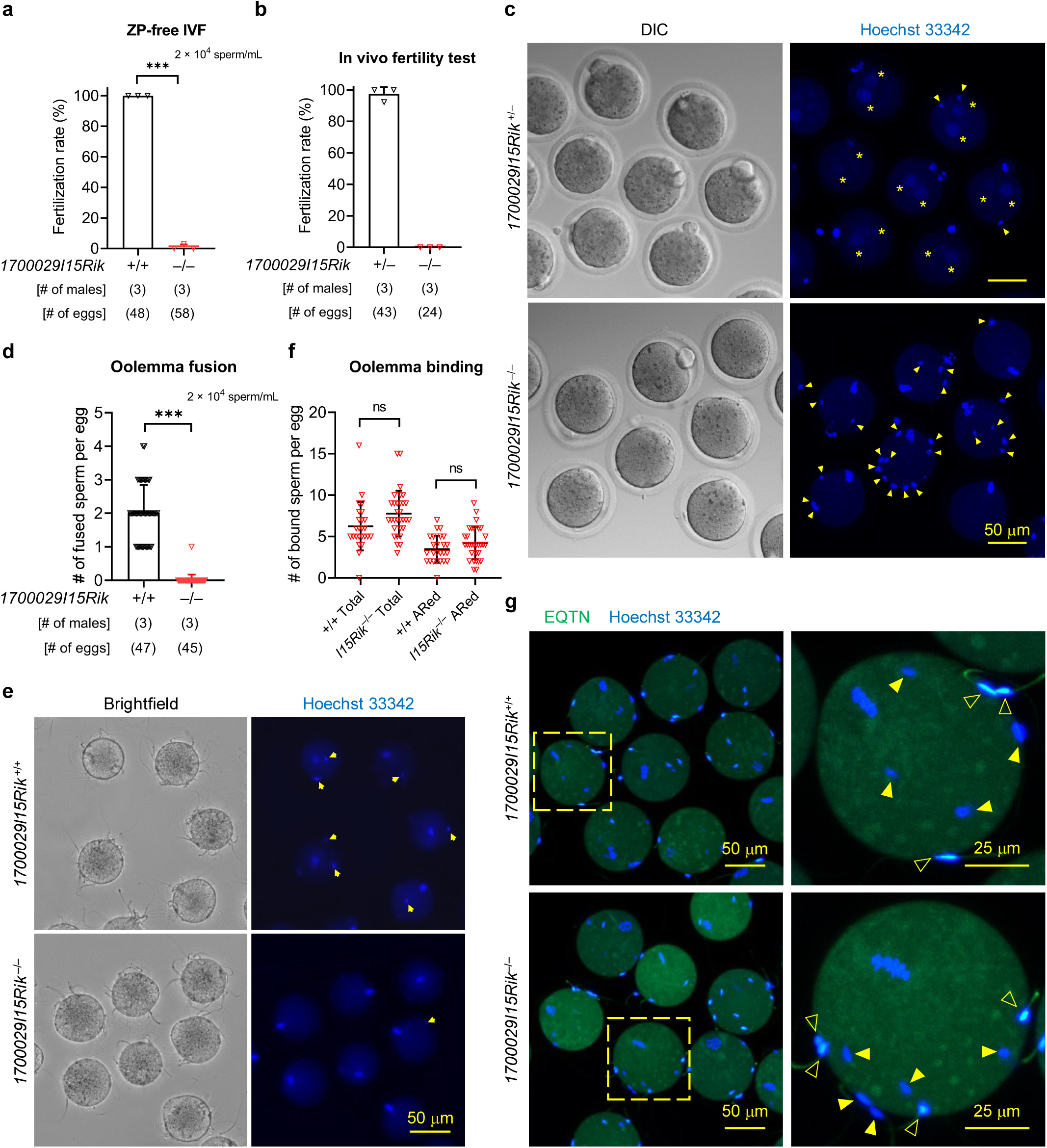
Infertility of *1700029I15Rik* knockout males is attributed to impaired sperm–egg fusion. **a**, In vitro fertilisation (IVF) with ZP-free wildtype eggs. **b – c**, In vivo fertility test of wildtype and knockout males. The eggs were harvested from superovulated B6D2F1 female mice that had copulated with wildtype or *1700029I15Rik* knockout males. Perivitelline sperm were visualised by Hoechst 33342 staining. **d – e**, Analysis of sperm-oolemma fusion using Hoechst 33342-preloaded ZP-free eggs. Yellow arrows indicate the fused sperm heads carrying the Hoechst dye transferred from the egg cytoplasm. **f – g**, Analysis of sperm-oolemma binding. Spermatozoa pre-incubated in the TYH medium were probed with an anti-EQTN antibody and fluorophore-conjugated secondary antibody to reveal the acrosomal status. The acrosome-intact and reacted spermatozoa are marked by solid and hollow arrowheads, respectively.

### 1700029I15Rik interacts with ER-resident proteins implicated in protein folding, N-glycosylation, and ER–Golgi trafficking

To identify the interacting proteins of 1700029I15Rik in testis, we next performed co-immunoprecipitation tandem mass spectrometry (co-IP/MS) analyses using antibody-crosslinked agarose resin (SI Appendix, Fig. S5a). As a result, 31 proteins (including 1700029I15Rik) were specifically and concurrently detected in the wildtype samples from at least two of the three biological replicates (Fig. 4a). Gene ontology (GO) and Kyoto Encyclopedia of Genes and Genomes (KEGG) analyses revealed that the 31 hits included proteins pivotal for N-glycosylation (e.g., DAD1, OSTC, and RPN2) and ER–Golgi vesicular trafficking (e.g., TMED4 and TMED10; Fig. 4a, b).

**Fig. 4.**
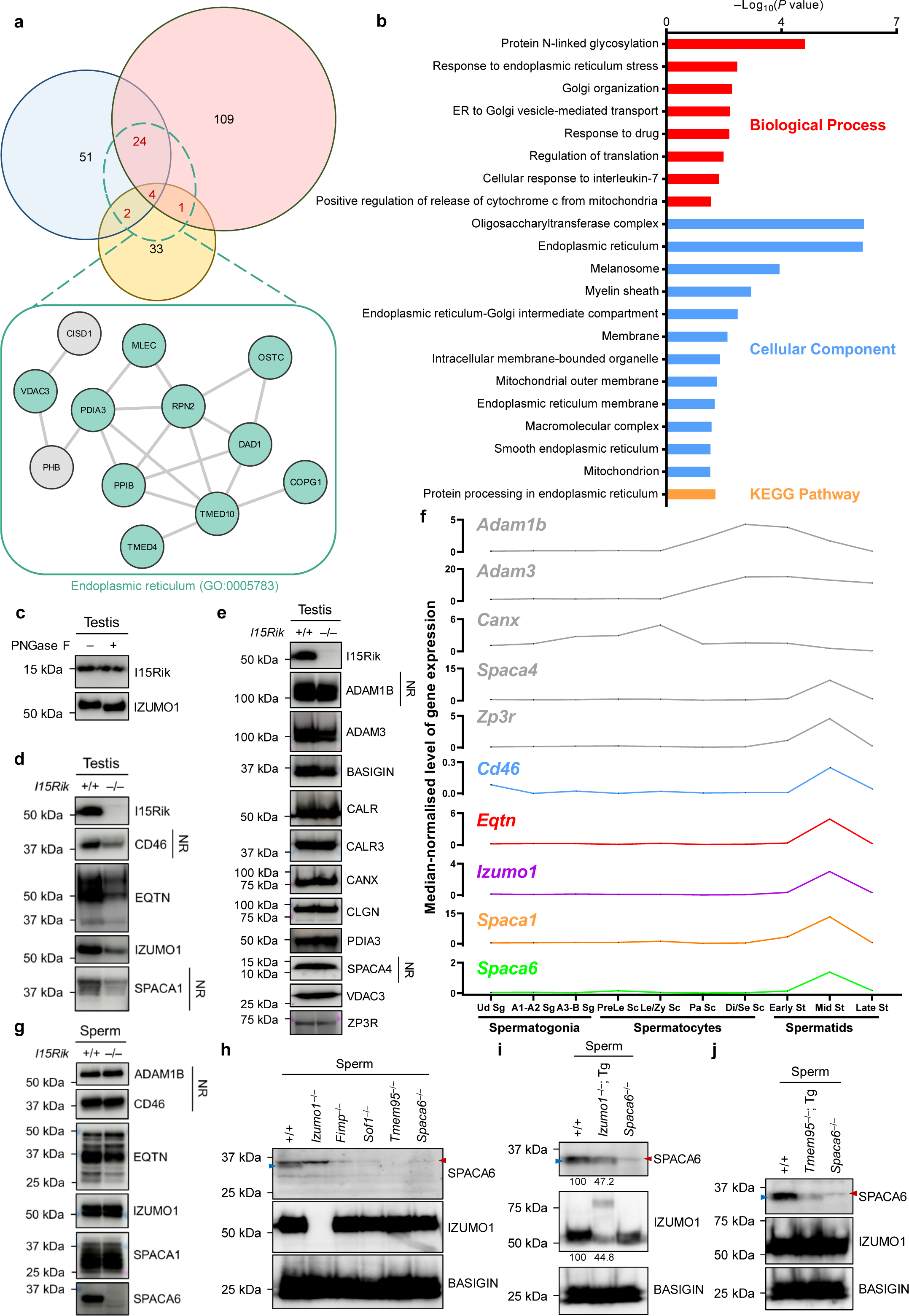
1700029I15Rik interacts with proteins involved in N-glycosylation and vesicular trafficking and underpins the assembly of acrosomal membrane proteins. **a**, Venn diagram and STRING analysis (29, 30) of concurrent proteins identified in three replicates of co-IP/MS experiments. Only the proteins specifically detected in the wildtype samples were analysed by STRING. **b**, GO and KEGG analyses of the highly associated interacting proteins. Functional annotations of interactome were conducted using the DAVID knowledgebase (31). Only the 31 proteins concurrently detected in at least two replicates of the co-IP/MS experiments were analysed. **c**, N-glycosylation analysis of testis 1700029I15Rik by PNGase F digestion. IZUMO1, a typical N-linked glycoprotein in the testis, was analysed as a positive control. **d – e**, Western blot analyses of various proteins in wildtype and knockout testes. Unless specified otherwise, all protein samples were processed under the reducing condition. NR, non-reducing. **f**, mRNA expression of *Adam1b*, *Adam3*, *Canx*, *Cd46*, *Eqtn*, *Izumo1*, *Spaca1*, *Spaca4*, and *Spaca6*, and *Zp3r* in the mouse spermatogenic cells based on a published scRNA-seq dataset (12). Ud Sg, undifferentiated spermatogonia; A1-A2 Sg, types A1 to A2 spermatogonia; A3-B Sg, types A3 to B spermatogonia; PreLe Sc, preleptotene spermatocytes; Le/Zy Sc, leptotene/zygotene spermatocytes; Pa Sc, pachytene spermatocytes; Di/Se Sc, diplotene/secondary spermatocytes; St, spermatids; Early St, early round spermatids; Mid St, mid-round spermatids; Late St, late round spermatids. **g**, Western blot analyses of various proteins in wildtype and knockout spermatozoa. **h**, Western blot analyses of the level of SPACA6 in *Izumo1*, *Fimp*, *Sof1*, *Tmem95*, and *Spaca6* knockout spermatozoa. Blue and red arrowheads mark the SPACA6 protein band and non-specific bands introduced by immunoglobulin G (IgG) contamination, respectively. **i**, Western blot analyses of the levels of SPACA6 and IZUMO1 in the sperm from wildtype, *Izumo1*^−/−^; Tg, and *Spaca6*^−/−^ males. Band intensities were measured by ImageJ and relative intensities were calculated with the wildtype bands set to 100. **j**, Western blot analyses of the levels of SPACA6 in the sperm from wildtype, *Tmem95*^−/−^; Tg, and *Spaca6*^−/−^ males.

Since 1700029I15Rik interacts with catalytic subunits of the oligosaccharyltransferase (OST) complex required for N-glycan biogenesis (e.g., DAD1 and OSTC), we next asked whether 1700029I15Rik itself undergoes N-linked glycosylation. By Peptide-N-Glycosidase F (PNGase F) digestion, we discovered that 1700029I15Rik did not undergo N-linked glycosylation in vivo or in vitro (Fig. 4c and SI Appendix, Fig. S6a, b). Thus, we suspected 1700029I15Rik plays a role in the N-glycan biogenesis.

To identify additional interactome of 1700029I15Rik with high confidence, another co-IP/MS analysis using non-crosslinked magnetic beads was incorporated, and the specificity of protein–protein interactions was scored based on the detected spectra (SI Appendix, Fig. S5b – d). Apart from the abovementioned interacting proteins, SPPL2C, PDIA3, and VDAC3 were found tightly associated with 1700029I15Rik (SI Appendix, Fig. S5d). SPPL2C is a signal peptide peptidase that selectively cleaves type-II transmembrane proteins (13), but it is dispensable for the male fertility (14). Interestingly, the level of 1700029I15Rik was dramatically decreased in *Sppl2c* knockout testis (13), suggesting SPPL2C is the upstream regulator underpinning the expression of 1700029I15Rik. PDIA3 is an ER-resident protein disulfide isomerase that modulates protein folding, formation and remodelling of disulfide bonds, and quality control of glycoproteins (15). It might be implicated in the sperm ZP binding and oolemma fusion, as revealed by previous inhibitory studies (16, 17).

### 1700029I15Rik ensures proper assembly of multiple transmembrane glycoproteins in the acrosome

Since our co-IP/MS results suggest potential involvement of 1700029I15Rik in protein processing and trafficking, we examined the expression levels of gamete fusion-related sperm proteins in *1700029I15Rik* knockout males. Western blot analyses indicated the amounts of acrosomal membrane glycoproteins, CD46, EQTN, IZUMO1, and SPACA1, were drastically reduced in the knockout testis (Fig. 4d and SI Appendix, Fig. S6c – e). Nonetheless, the levels of sperm head plasma membrane proteins (e.g., ADAM1B and ADAM3), ER chaperones (e.g., CALR, CANX, and PDIA3), SPACA4, a GPI-anchored protein localised to the acrosomal membrane, and ZP3R, a soluble protein expressed in the acrosomal matrix, were comparable in wildtype and knockout testis (Fig. 4e). The downregulated acrosomal membrane proteins are highly expressed in the mid-round spermatids, whereas the plasma membrane proteins show peaked expression in the spermatocytes (Fig. 4f).

Remarkably, we further discovered that the proteins downregulated in *1700029I15Rik*^−/−^ testis exhibited normal expression levels in the knockout sperm (Fig. 4g). The localisations of EQTN, IZUMO1, and SPACA1 were also normal in the knockout spermatozoa (SI Appendix, Fig. S6f). SPACA6, however, was absent in the *1700029I15Rik* knockout spermatozoa, thus providing a plausible explanation for the defective oolemma fusion ability (Fig. 4g and SI Appendix, Fig. S6g).

A previous study has indicated SPACA6 is absent in *Izumo1*, *Dcst1*, and *Dcst2* knockout sperm (8). Here, we found that SPACA6 was also missing in *Tmem95*, *Fimp*, and *Sof1* knockout sperm (Fig. 4h). In the spermatozoa from *Izumo1*^−/−^; Tg males, the level of transgenic IZUMO1 was lower than the endogenous level of IZUMO1, and the transgene did not fully restore SPACA6 to its endogenous level (Fig. 4i). Intriguingly, a similar phenomenon was observed in the sperm from *Tmem95*^−/−^; Tg males (Fig. 4j). We thus hypothesised that the presence and abundance of SPACA6 is dependent upon that of the other fusion-related proteins. RT-PCR revealed that the mRNA expression of *Spaca6* was normal in *Tmem95*^−/−^ and *Tmem95*^−/−^; Tg testis (SI Appendix, Fig. S6h), indicating the abundance of SPACA6 was influenced by the other factors at the protein level. Additionally, the amount of 1700029I15Rik was normal in *Izumo1* and *Spaca6* knockout males (SI Appendix, Fig. S6i, j). Collectively, these data indicate the depletion of 1700029I15Rik causes aberrant assembly of fusion-related acrosomal membrane proteins and the missing of SPACA6.

### 1700029I15Rik is an ER-expressing protein underpinning proper folding of acrosomal membrane proteins

To investigate the molecular function of 1700029I15Rik, we carried out in vitro analyses using the HEK293T cells. The plasmid encoding C-terminal 3×FLAG-tagged 1700029I15Rik was used because it exhibited normal functioning in vivo, reflected by the successful restoration of the fertility in the knockout males (Fig. 2d and SI Appendix, Fig. S6k). C-terminal tagged 1700029I29Rik displayed a normal interaction with SPPL2C in vitro (SI Appendix, Fig. S6l), further validating the rationale for using this construct for in vitro analyses.

In line with the discovery that 1700029I15Rik interacts with multiple ER proteins, we found this protein was predominantly co-localised with KEDL, an ER-resident protein, in the HEK293T cells (Fig. 5a). To investigate how 1700029I15Rik would facilitate protein processing, we co-transfected HEK293T cells with plasmids encoding 1700029I15Rik and other fusion-related proteins. Murine TMEM95 contains two N-linked glycosylation sites, N36 and N118. The presence of 1700029I15Rik significantly enhanced the expression amount of the non-glycosylated, but not the fully glycosylated TMEM95 (SI Appendix, Fig. S7a and Fig. 5b, c). Proteinase K treatment revealed no significant change in the surface expression of TMEM95 with or without the presence of 1700029I15Rik (Fig. 5d). These findings imply that 1700029I15Rik might be involved in the processing of glycoproteins prior to the N-glycan attachment. Distinctively, 1700029I15Rik did not improve the expression of IZUMO1 or SPACA6, that were efficiently and extensively glycosylated in the HEK293T cells (SI Appendix, Fig. S7b, c).

**Fig. 5.**
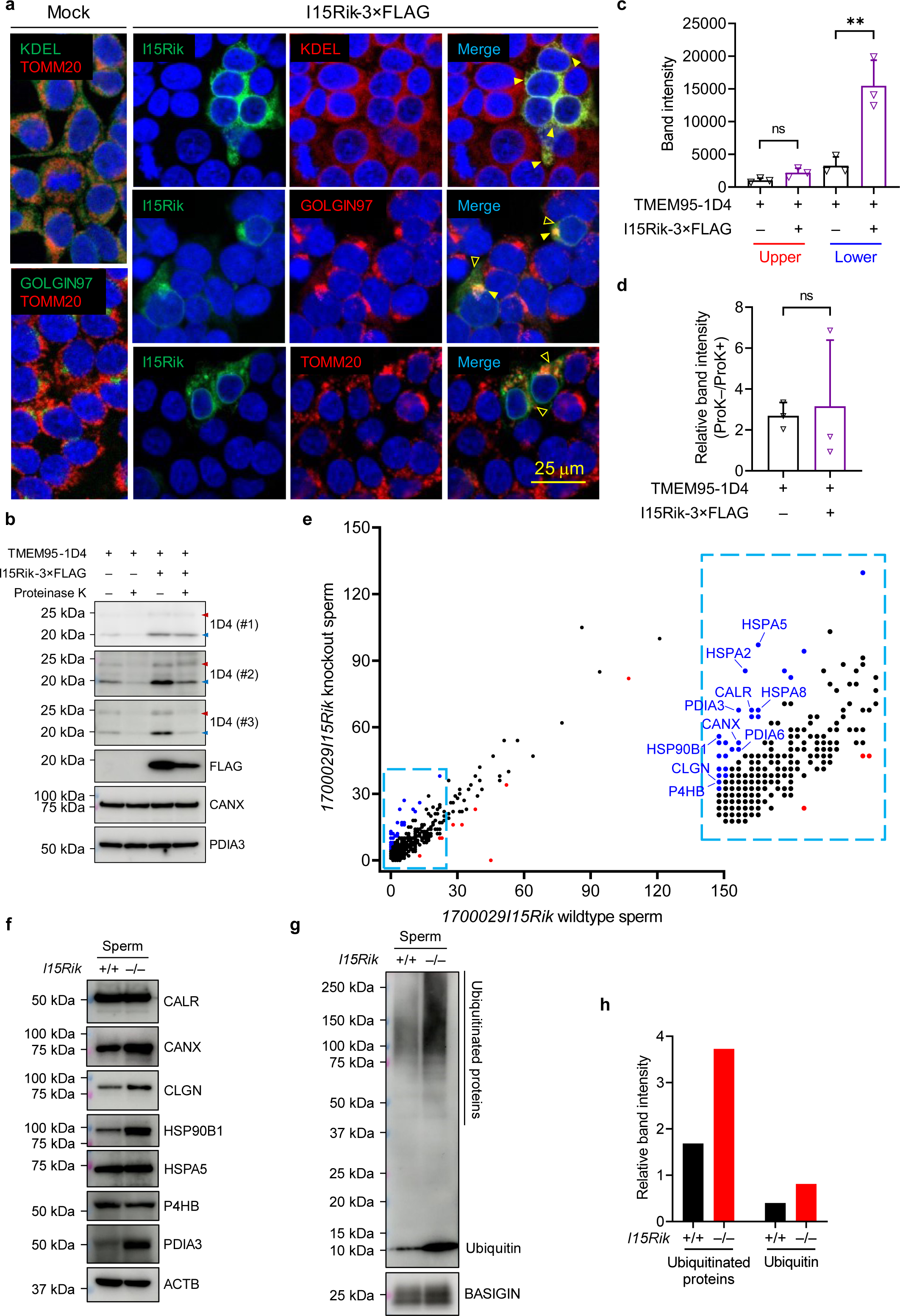
Ablation of 1700029I15Rik results in upregulation of the ERAD machinery and elevation of ubiquitinated proteins in the sperm. **a**, Immunostaining of HEK293T cells transiently expressing 1700029I15Rik. Solid arrowheads indicate co-localisation of 1700029I15Rik and other organelle markers, whereas hollow arrowheads indicate the proteins are not co-localised. The cell nuclei were visualised by Hoechst 33342. **b**, Co-expression of 1700029I15Rik and TMEM95 in HEK293T cells. To examine the surface expression, HEK293T cells were treated with Proteinase K to shave off the surface proteins. CANX and PDIA3 were analysed in parallel as loading controls. The red and blue arrowheads indicate glycosylated and non-glycosylated TMEM95, respectively. **c**, Analysis of total expression of TMEM95 with or without the presence of 1700029I15Rik. The band intensities were measured by ImageJ. **d**, Analysis of the surface expression of TMEM95 with or without the presence of 1700029I15Rik. Relative band intensities were calculated by dividing the untreated with the Proteinase K (ProK)-treated group. **e**, MS analysis of 1700029I15Rik wildtype and knockout sperm proteome. Blue and red dots indicate proteins upregulated and downregulated in the knockout spermatozoa, respectively. **f**, Western blot analyses of the levels of chaperone proteins in the wildtype and knockout spermatozoa. ACTB was analysed as a loading control. **g**, Western blot analyses of the amounts of ubiquitin and ubiquitinated proteins in the wildtype and knockout spermatozoa. **h**, Relative band intensities of ubiquitin and ubiquitinated proteins in the wildtype and knockout spermatozoa. The band intensities were measured and relative intensities of ubiquitin or ubiquitinated proteins relative to BASIGIN were calculated by ImageJ.

By MS, we further discovered that a multitude of proteins were upregulated in the knockout sperm (Fig. 5e). GO and STRING analyses indicated most of these upregulated proteins are ER chaperones that exhibit unfolded or misfolded protein binding ability (e.g., CANX, CALR, and HSP90B1) or peptide disulfide oxidoreductase activity (e.g., P4HB, PDIA3, and PDIA6; SI Appendix, Fig. S7d – f). By western blot, we confirmed the amounts of CANX, CLGN, HSP90B1, and PDIA3 were higher in the *1700029I15Rik* knockout spermatozoa, whereas the levels of CALR, HSPA5, and P4HB remained unchanged (Fig. 5f). We further observed an increase of ubiquitin and ubiquitinated proteins in the knockout spermatozoa, indicating the ubiquitin-dependent ER-associated degradation (ERAD) is upregulated (Fig. 5g, h). Notably, in the HEK293T cells, the amounts of CANX and PDIA3 were not changed in response to the absence of 1700029I15Rik (Fig. 5b), indicating the in vitro co-expression system does not fully recapitulate the in vivo situation. Overall, the in vitro and proteomic studies suggest potential involvement of 1700029I15Rik in governing proper assembly of the nascent acrosomal membrane proteins during spermiogenesis.

## Discussion

Nascent polypeptides are translocated into the ER and fold into their native conformations. To maintain protein homeostasis, unfolded or misfolded proteins are eliminated by sophisticated quality control systems (18). In somatic cells, lectin-like ER chaperones calnexin (CANX) and Calreticulin (CALR) target misfolded proteins for degradation via the ubiquitin-dependent ERAD pathway (18, 19). In the male germline, Calmegin (CLGN) and Calreticulin-3 (CALR3), the testis-specific paralogues of CANX and CALR, respectively, regulate the biosynthesis of sperm plasma membrane proteins (e.g., ADAM1B and ADAM3) required for pre-fertilisation events (20–22). Nevertheless, ablation of CLGN or CALR3 does not affect sperm– egg fusion. Thus, the machinery facilitating the biogenesis of gamete fusion-related proteins remains elusive (22, 23).

As revealed by co-IP/MS, 1700029I15Rik interacts with ER proteins involved in disulfide bond remodelling, N-linked glycosylation, and ER–Golgi vesicular trafficking (Fig. 4a, b and SI Appendix, Fig. S5). In *1700029I15Rik* knockout males, a number of acrosomal membrane proteins are downregulated in the testis, while the ubiquitinated and ERAD-associated proteins are accumulated in the sperm (Fig. 4d and Fig. 5f, g). These findings raise the possibilities that 1700029I15Rik potentiates the proper folding, attachment of N-glycans, or intracellular transport of the nascent acrosomal membrane proteins by recruiting corresponding ER chaperones. However, IZUMO1 and other putative substrates remain targeted to the acrosomal membrane (Fig. 4g and SI Appendix, Fig. S6f), implying 1700029I15Rik might not be the sole chaperone processing the acrosomal membrane proteins.

In addition to the aforementioned possibilities, we propose an alternative hypothetical model to be interrogated in the future research: 1700029I15Rik might enable rapid formation of folding intermediates that cannot be targeted by the quality control machinery. In the absence of 1700029I15Rik, nascent slow-folding proteins are directed to the ERAD pathway, whereas fast-folding proteins are escaped from targeted destruction (SI Appendix, Fig. S7g). It is generally accepted that small proteins less than 100 residues fold rapidly through two-state kinetics, whereas larger proteins populate folding intermediates before transforming to their native conformations (24–26). The known gamete fusion-related sperm proteins vary greatly in their sizes and the numbers of disulfide bridges. If this hypothetical model stands, different fusion-related factors might exhibit varied degrees of reduction in response to the depletion of 1700029I15Rik.

According to the scRNA-seq analysis (12), proteins destined for the plasma membrane and the acrosomal membrane exhibit striking differences in their timing of mRNA expression. For example, the plasma membrane proteins, ADAM1B and ADAM3, and acrosomal membrane proteins, IZUMO1 and SPACA1, show peaked mRNA expression in the late spermatocytes and mid-round spermatids, respectively (SI Appendix, Fig. S1b and Fig. 4f). Given that only the acrosomal membrane proteins have so far been discovered downregulated in the *1700029I15Rik* knockout testis, it is tempting to speculate that the biosynthesis of plasma membrane and acrosomal membrane proteins is regulated in a spatiotemporal manner. However, whether the substrates of 1700029I15Rik are restricted to the acrosomal membrane proteins needs to be investigated.

SPACA6 is absent in *1700029I15Rik* knockout spermatozoa. Remarkably, it is also missing in the knockout sperm lacking any of the known fusion-related factors [Fig. 4h, (8)]. The level of SPACA6 decreases with the reduction of IZUMO1 and TMEM95 in the transgenic males (Fig. 4i, j), suggesting IZUMO1, TMEM95, and perhaps other fusion-related proteins assemble into a complex on the acrosomal membrane, where SPACA6 subsequently binds to. To untangle the detailed architecture of this putative protein complex, antibodies or alternative immunodetection approaches shall be developed and employed in future analyses.

The functional specialisation of the male germline and a lack of culture systems that reconstitute the germ cell differentiation impede us from fully recapitulating the 1700029I15Rik-mediated protein processing pathway in vitro. Prominently, this work unravels 1700029I15Rik as a key regulator in the biosynthesis of sperm acrosomal membrane proteins. This pathway acts parallelly to the CLGN/CALR3-mediated processing of sperm plasma membrane proteins. Our discovery emphasises the importance of proper biosynthesis and correct assembly of acrosomal membrane proteins, the failure of which poses far-reaching adverse consequences on the fertilisation process. This knowledge potentially contributes to developing a novel contraceptive measure involving molecular perturbations in the biogenesis of gamete fusion-required acrosomal proteins.

## Acknowledgements (168 words)

The authors thank Natsuki Furuta, Saki Nishioka, and the Biotechnology Research and Development (nonprofit organization) for technical assistance; Akinori Ninomiya and the Core Instrumentation Facility for assistance in MS analyses; the members at Department of Experimental Genome Research for critical discussion. This work was supported by the Ministry of Education, Culture, Sports, Science, and Technology (MEXT)/Japan Society for the Promotion of Science (JSPS) KAKENHI grants (JP18K16735 and JP22K15103 to Y.L., JP18K14612 and JP20H03172 to T.N., and JP19H05750 and JP21H05033 to M.I.), the Takeda Science Foundation grants to T.N. and M.I., the Japan Science and Technology Agency (JST) PRESTO grant JPMJPR2148 to T.N., the Nakajima Foundation to T.N., the Eunice Kennedy Shriver National Institute of Child Health and Human Development grants K99HD104924 to S.T., and P01HD087157 and R01HD088412 to M.I., the Swedish Research Council grant 2018-03426 to H.S., and the Bill & Melinda Gates Foundation grant INV-001902 to M.I. The funders had no role in study design, data collection and analysis, decision to publish or preparation of the manuscript.

## SI Appendix

### SI Materials and Methods

#### Animals

C57BL/6J, B6D2F1, ICR and SWISS mice were purchased from Japan SLC, Inc. (Shizuoka, Japan) or Janvier Labs (Le Genest Saint Isle, France) and maintained under cycles of 12 h light and 12 h darkness with *ad libitum* feeding. All animal experiments were approved by the Animal Care and Use Committee of the Research Institute for Microbial Diseases, Osaka University (#Biken-AP-H30-01) and the University of Gothenburg (#1316/18), and were conducted in compliance with all guidelines and regulations. Mice were euthanised by cervical dislocation following anaesthesia.

Frozen spermatozoa from *1700029I15Rik* heterozygous knockout males (B6D2-1700029I15Rik<em1OSB>, RBRC#11049, CARD#2956), and *1700029I15Rik* transgenic (Tg) males (B6D2-Tg(Clgn-1700029I15Rik/PA)1Osb, RBRC#11470, CARD#3114) are available at the Riken BioResource Center (RIKEN BRC; web.brc.riken.jp/en) and the Center for Animal Resources and Development, Kumamoto University (CARD R-BASE; cardb.cc.kumamoto-u.ac.jp/transgenic).

#### Antibodies and plasmids

Antibodies used in this study are listed in <u>SI Appendix,</u> Table S2. In the present study, polyclonal antibodies against the aa 81 – 99 of mouse 1700029I15Rik (QQRKRDGPNMADYYYDVNL) and the aa 75 – 93 of mouse SPACA6 (EGAFEDLKDMKINYDERSYL) were raised and produced in rabbit by Sigma (Sigma-Aldrich, St. Louis, MO).

The open reading frames (ORFs) of *1700029I15Rik*, *Izumo1*, *Spaca6*, *Sppl2c*, and *Tmem95* were cloned and amplified from mouse testis cDNA by PCR and inserted into the multiple cloning site of the pCAG1.1 vector (Addgene; Plasmid #173685).

#### In silico analysis of gene expression in spermatogenic cells

The mRNA expression of mouse *1700029I15Rik*, *Adam1b*, *Adam3*, *Canx*, *Cd46*, *Dcst1*, *Dcst2*, *Eqtn*, *Fimp*, *Izumo1*, *Sof1*, *Spaca1*, *Spaca4*, *Spaca6*, *Tmem95*, and *Zp3r* in the male germ cells at different spermatogenic stages was evaluated based on the scRNA-seq dataset published by Hermann et al. (1).

#### Reverse transcription polymerase chain reaction (RT-PCR)

Total RNA was extracted from various tissues and organs of C57BL/6J mice by TRIzol (ThermoFisher, Waltham, MA) and reverse transcribed to cDNA using SuperScript III First Strand Synthesis Kit (ThermoFisher, Waltham, MA) as per manufacturer’s instruction. The mRNA expression was then analysed by PCR using the KOD Fx Neo DNA polymerase (Toyobo, Tokyo, Japan). Sequences of the primers for RT-PCR are listed in the SI Appendix, Table S3.

#### Generation of *170029I15Rik* knockout, mutant, and transgenic mouse lines

*1700029I15Rik* knockout mice were generated by the CRIPSR/Cas9-based gene editing technology as previously described (2). Two sgRNAs targeting the first coding exon and the 3’ UTR of *1700029I15Rik* were designed to remove the entire coding sequence. Wildtype two-pronuclear (2PN) zygotes were collected from the B6D2F1 female mice that had been superovulated by peritoneal injection of CARD HyperOva (Kyudo Co., Saga, Japan) and human chorionic gonadotropin (hCG; ASKA Pharmaceutical Co. Ltd., Tokyo, Japan) and paired with B6D2F1 males. A ribonucleoprotein complex of CRISPR RNA (crRNA), trans-activating crRNA (tracrRNA) and Cas9 protein was introduced into the 2PN eggs using a NEPA21 super electroporator (NEPA GENE, Chiba, Japan). The electroporated zygotes were cultured in the potassium simplex optimisation medium (KSOM) to two-cell stage and transplanted into the ampullary segment of the oviducts of 0.5-d pseudopregnant ICR females. The founder generation was obtained by natural delivery or Cesarean section and genotyped by PCR using primers (Fw and Rv) targeting the introns upstream and downstream of the coding region. The mutant allele was subsequently verified by Sanger sequencing.

Similarly, the *1700029I15Rik* mutant mouse line was generated using two sgRNAs targeting the upstream and downstream introns of the first coding exon. Mouse zygotes were obtained from mating between C57BL/6J females and males. Cas9 protein (Sigma-Aldrich, St. Louis, MO) and sgRNAs were mixed to form ribonucleoprotein complexes and co-injected into the zygotes in the M2 medium. Microinjected zygotes were allowed to recover for 2 – 4 h in the KSOM medium in a humidified CO_2_ incubator at 37°C and were then transferred into pseudopregnant SWISS female mice. Likewise, the founders were genotyped using primers (Fw’ and Rv’) flanking the mutation site and the PCR products were subjected to Sanger sequencing. The founder mice were crossed with C57BL/6J mice to avoid potential off-target mutations.

For producing the transgenic mouse line, the ORF of *1700029I15Rik* was cloned from mouse testis cDNA by PCR and inserted into a plasmid encoding a *Clgn* promoter and a rabbit beta globin polyadenylation (polyA) signal. Linearised plasmids were microinjected into the pronuclei of zygotes obtained from mating between homozygous knockout females and heterozygous knockout males. The injected zygotes were cultured in the KSOM medium until the two-cell stage and transplanted into the oviductal ampulla of 0.5 d pseudopregnant ICR females. Founder animals were obtained by natural delivery or Cesarean section after 19 d of pregnancy. The transgene was identified by PCR using primers (Tg_Fw and Tg_Rv) targeting the *Clgn* promoter and the polyA signal.

Sequences of the primers used for genotyping of the knockout, mutant, and transgenic mice are enumerated in the SI Appendix, Table S3.

#### Fertility tests

Upon sexual maturation, homozygous knockout male mice or knockout Tg males were caged individually with three 6-week-old wildtype B6D2F1/J female mice for eight weeks. During this period, vaginal plugs was examined as an indicator of successful copulation and the number of pups was counted at birth. Three knockout males were tested to meet the requirements for statistical validity. The fecundity of three wildtype B6D2F1/J males was tested in parallel as positive controls. After the eight-week mating period, the male mice were withdrawn from the cages and the females were kept for another three weeks to allow their final litters being delivered.

For the fertility tests of 1700029I15Rik mutant mice, one male was housed with one female mouse consecutively for six months. The number of pups were recorded at birth.

#### In vitro fertilisation

Cauda epididymal spermatozoa from *1700029I15Rik* heterozygous and homozygous knockout males were dispersed in the Toyoda, Yokoyama, Hoshi (TYH) medium and pre-incubated for 2 h at 37°C, 5% CO_2_ to induce capacitation. Cumulus-oocyte complexes (COCs) were extracted from the oviductal ampulla of superovulated B6D2F1 female mice, and were treated with 330 μg/mL of hyaluronidase (Wako, Osaka, Japan) to remove the cumulus cells, or 1 mg/mL of collagenase (Sigma-Aldrich, St. Louis, MO) to remove the ZP. Cumulus-intact and cumulus-free oocytes were incubated with spermatozoa at a concentration of 2 × 10^5^ sperm/mL in 100 µL TYH medium drops, whereas ZP-free oocytes were inseminated at 2 × 10^4^ sperm/mL. After 6 h of insemination, fertilisation success was determined by formation of two pronuclei.

#### In vivo fertility test

B6D2F1 female mice were superovulated by peritoneal injection of PMSG and hCG and paired with wildtype or *1700029I15Rik*^−/−^ males. Oviducts were isolated from the female mice about 6 h after the formation of copulation plugs. COCs were extracted from the oviductal ampulla, released in the KSOM medium drops, and treated with 330 µg/mL of hyaluronidase (Wako, Osaka, Japan) to remove the cumulus cells. The eggs were then transferred to FHM medium drops on glass bottom culture dishes and movies were captured under a Keyence BZ-X810 microscope (Keyence, Osaka, Japan). The eggs were then fixed in 0.25% glutaraldehyde in FHM medium on ice for 15 min and stained with Hoechst 33342 (1:1000) to visualise the pronuclei and perivitelline sperm. Fluorescence images were captured under a Nikon Eclipse Ti microscope equipped with a Nikon C2 confocal module (Nikon, Tokyo, Japan).

#### Sperm–egg binding and fusion assay

For analysing sperm binding to the oolemma, cauda epididymal spermatozoa were dispersed in TYH medium and pre-incubated at 37°C, 5% CO_2_ for 2 h. After the pre-incubation, the spermatozoa were incubated for another 30 min in a TYH medium drop containing anti-EQTN antibody and Alexa Fluor 488 anti-mouse secondary antibody at a density of 2 × 10^6^ sperm/mL. In the meantime, COCs were extracted from superovulated females and treated with 1 mg/mL of collagenase to remove both the cumulus cells and the ZP. ZP-free eggs were incubated with antibody-probed spermatozoa at a final concentration of 2 × 10^4^ sperm/mL. After insemination for 30 min, the sperm–egg complexes were fixed in 0.25% glutaraldehyde in FHM medium on ice for 15 min and stained with Hoechst 33342 (1:1000) to visualise the bound sperm. The total number of bound sperm and the number of acrosome-reacted sperm bound to the oolemma was calculated under a Keyence BZ-X810 microscope (Keyence, Osaka, Japan). Z-stack images of the sperm–egg complexes were captured under a Nikon Eclipse Ti microscope equipped with a Nikon C2 confocal module (Nikon, Tokyo, Japan).

#### Sperm ZP binding assay

Cauda epididymal sperm were pre-incubated in the TYH medium for 2 h at 37°C, 5% CO_2_. The eggs harvested from B6D2F1 female mice were denuded by treating with 330 µg/mL hyaluronidase at 37°C for 5 min. The pre-incubated sperm were added to fresh TYH medium drops containing cumulus-free eggs at a sperm density of 2 × 10^5^ sperm/mL and incubated at 37°C, 5% CO_2_ for 30 min. The sperm–egg complexes were fixed in 0.25% glutaraldehyde for 15 min on ice, rinsed in fresh FHM medium drops for three times, and examined under an Olympus BX-53 differential interference contrast microscope equipped with an Olympus DP74 colour camera (Olympus, Tokyo, Japan).

#### Cell-egg binding assay

HEK293T cells were transfected with plasmids encoding 3×FLAG-tagged 1700029I15Rik or mCherry-tagged IZUMO1 by the calcium phosphate–DNA co-precipitation method. Mock transfection was carried out using a blank vector bearing a CAG promoter and a polyA signal. Approximately 18 hours after transfection, cells were detached by repeatedly pipetting, washed in PBS, and resuspended in the TYH medium. HEK293T cells were mixed with ZP-free eggs in fresh TYH medium drops and incubated at 37°C, 5% CO_2_ for 30 min. The eggs were washed briefly to remove loosely bound cells and observed under a Keyence microscope.

#### Histological analyses of testes and epididymides

Testes and epididymides were fixed in Bouin’s solution (Polysciences, Inc., Warrington, PA), embedded in paraffin wax, and sectioned at a thickness of 5 – 10 μm on a Microm HM325 microtome (Microm, Walldorf, Germany). The paraffin sections were stained with periodic acid (Nacalai Tesque, Kyoto, Japan) and Schiff’s reagent (Wako, Osaka, Japan), followed by counterstaining with Mayer’s haematoxylin solution (Wako, Osaka, Japan). Alternatively, the testis sections were stained with haematoxylin and eosin.

#### Sperm morphology and motility analyses

Spermatozoa were extracted from cauda epididymides and dispersed in PBS or TYH medium. Sperm morphology was observed under an Olympus BX-53 differential interference contrast microscope equipped with an Olympus DP74 colour camera (Olympus, Tokyo, Japan). Sperm motility was measured by the CEROS II sperm analysis system (Hamilton Thorne Biosciences, Beverly, MA) at 10 min and 2 h of incubation in the TYH medium.

#### Immunostaining of spermatocyte chromosome spreads

Testicular germ cells were extracted from the wildtype and mutant testes, washed in PBS, centrifuged, and resuspended in hypotonic buffer [30 mM Tris (pH 7.5), 17 mM trisodium citrate, 5 mM EDTA, and 50 mM sucrose], followed by centrifugation and resuspension in 100 mM sucrose. The cell suspensions were then smeared on microscope slides in a same volume of fixation buffer (1% paraformaldehyde and 0.1% Triton X-100), fixed at room temperature for 3 h, and air dried. For immunostaining, the testicular germ cells were incubated with primary antibodies in PBS containing 5% BSA for 2 h, followed by fluorophore-conjugated secondary antibodies for 1 h at room temperature. The slides were washed with PBS and mounted with VECTASHIELD Antifade Mounting Medium with DAPI (Vector Laboratories, Burlingame, CA) prior to imaging.

#### Western blot

Mouse testes and epididymal sperm were lysed in 1% Triton X-100 lysis buffer, radioimmunoprecipitation (RIPA) buffer, or sodium dodecyl sulfate (SDS) sample buffer [66.7 mM Tris-HCl (pH 6.8), 2% (w/v) SDS, 0.003% (w/v) bromophenol blue, 10% Glycerol, and 5% β-mercaptoethanol in distilled water]. Protein lysates were resolved by sodium dodecyl sulfate-polyacrylamide gel electrophoresis (SDS-PAGE) and transferred to polyvinylidene fluoride (PVDF) membranes using the Trans-Blot Turbo system (BioRad, Munich, Germany). After blocking with 10% Difco™ skim milk (BD Bioscience, Sparks, MD) in Tris-Buffered Saline containing 0.1% Tween 20 (TBST; Nacalai Tesque, Kyoto, Japan), the PVDF membranes were incubated with primary antibodies for 3 h at room temperature or overnight at 4°C. The membranes were then probed with horseradish peroxidase (HRP)-conjugated secondary antibodies (Jackson ImmunoResearch Laboratories, West Grove, PA) for 1 h at room temperature and immersed in Amersham™ ECL™ Western Blotting Reagent Pack (Cytiva, Tokyo, Japan) for 15 s. Chemiluminescence was detected by Amersham™ ImageQuant™ 800 (Cytiva, Tokyo, Japan).

#### Immunocytochemistry and immunohistochemistry

Spermatozoa from cauda epididymides were air dried on microscope slides and fixed in 4% PFA in phosphate buffered saline (PBS) for 30 min at room temperature. The fixed spermatozoa were permeabilised with 0.1% Triton X-100 in PBS for 10 min at room temperature and blocked with 3% bovine serum albumin and 10% goat serum in PBS for 1 h at room temperature. The samples were then incubated with primary antibodies for 3 h at room temperature or overnight at 4°C. After three washes in 0.1% Triton X-100 in PBS, the spermatozoa were further incubated with fluorophore-conjugated secondary antibodies for 1 h at room temperature and with 1 μg/mL Hoechst 33342 (ThermoFisher, Waltham, MA) for 30 min at room temperature. The spermatozoa were mounted with VECTASHIELD Antifade Mounting Medium (Vector Laboratories, Burlingame, CA) prior to imaging. Fluorescence images were captured with an Olympus BX[53 microscope (Olympus, Tokyo, Japan) or a Nikon Eclipse Ti microscope equipped with a Nikon C2 confocal module (Nikon, Tokyo, Japan).

Testes were fixed in 4% paraformaldehyde and infused with 15% and 30% sucrose in PBS. The testes were then embedded in the OCT medium and snap-frozen in liquid nitrogen. Testis blocks were sectioned at a thickness of 5 – 10 μm on a cryostat and the sections were dried on adhesive microscope slides. The testis sections were then processed and imaged in the same manner as sperm immunostaining.

#### Triton X-114 fractionation of testis and sperm proteins

Testes were homogenised on ice in 1% Triton X-114 in PBS containing 1% (v/v) protease inhibitor cocktail (Nacalai Tesque, Kyoto, Japan) using a Dounce homogeniser. Cauda epididymal sperm were lysed in the same buffer at 4°C for 1 h with occasional vortexing. The insoluble cell debris were removed after centrifugation at 15,000 × g, 4°C for 30 min. The cleared testis and sperm lysates were then incubated at 37°C for 15 min and centrifuged at 15,000 × g, room temperature for 30 min. The upper aqueous phase was transferred to fresh conical tubes. To remove impurities from both fractions, the aqueous phase was mixed with Triton X-114 at a dilution of 1:100 and the detergent phase was mixed with 100 volumes of PBS. The solutions were fractionated again by incubation at 37°C and centrifugation at 15,000 × g, room temperature; the purification was repeated for three times. The purified detergent phase was diluted with PBS up to the same volume with the aqueous phase. An equal volume of both fractions was denatured in the SDS sample buffer at 95°C and subjected to SDS-PAGE and western blot analyses.

#### Co-immunoprecipitation

Testes were homogenised on ice in 1% Triton X-100 in PBS containing 1% (v/v) protease inhibitor cocktail (Nacalai Tesque, Kyoto, Japan) using a Dounce homogeniser. Co-IP was performed using the Invitrogen™ Dynabeads™ Magnetic Beads or Pierce™ Crosslink IP Kit (ThermoFisher, Waltham, MA) in accordance with the manufacturer’s instructions.

#### Silver staining

The protein samples were mixed with the SDS sample buffer, boiled at 95°C for 10 min, and resolved by SDS-PAGE. The gel was stained using Sil-Best Stain One (Nacalai Tesque, Kyoto, Japan) or Sil-Best Stain-Neo (Nacalai Tesque, Kyoto, Japan) according to the manufacturer’s instructions.

#### Mass spectrometry

Protein samples immunoprecipitated from testis lysates or extracted from mature spermatozoa were processed and the resultant protein peptides were subjected to nanocapillary reversed-phase liquid chromatography (LC)-MS/MS analysis using a C18 column (10 cm × 75 μm, 1.9 µm, Bruker Daltonics) on a nanoLC system (Bruker Daltonics, Billerica, MA) connected to a timsTOF Pro mass spectrometer (Bruker Daltonics, Billerica, MA) and a nano-electrospray ion source (CaptiveSpray; Bruker Daltonics, Billerica, MA). The resulting data were processed using DataAnalysis (Bruker Daltonics, Billerica, MA), and proteins were identified using MASCOT Sever (Matrix Science, Tokyo, Japan) compared with the SwissProt database. Quantitative value and fold exchange were calculated by Scaffold 5 software (Proteome Software, Portland, OR).

#### Cultured cell transfection and treatment

HEK293T cells were cultured in Dulbecco’s Modified Eagle Medium (DMEM) complete medium [Gibco™ DMEM (Thermofisher, Waltham, MA) supplemented with 10% foetal bovine serum (Sigma-Aldrich, St. Louis, MO) and 1% Gibco™ penicillin/streptomycin (Thermofisher, Waltham, MA)] at 37°C, 5% CO_2_. HEK293T cells were transfected by the calcium phosphate–DNA co-precipitation method. Cells were harvested by resuspending in PBS and lysed in 1% Triton X-100 lysis buffer at 4°C for 2 h with occasional vortexing.

To trim off the surface proteins, live cells were incubated in 0.1 mg/mL of Proteinase K in the DMEM complete medium at 37°C, 5% CO_2_ for 20 min. The treated cells were washed in PBS for three times and were lysed as described above.

#### Statistical analyses

Data are presented as mean values and error bars indicate standard deviation (SD). Experimental groups were analysed statistically using an unpaired two-tailed Student’s *t*-test. *P* values less than 0.05 were considered statistically significant (*, *P* < 0.05; **, *P* < 0.01; ***, *P* < 0.001).

#### Data availability

The raw data of MS analyses have been deposited to the ProteomeXchange Consortium via the PRIDE partner repository (3) with the dataset identifier PXD033246. The proteomics data are accessible with the username and password <u>Tgv6q4Yu</u> via the PRIDE database during the peer review process. All other data are included in the main text and the SI Appendix.

### SI Tables

**Table S1.** Discrepancies of 1700029I15Rik topology predicted by multiple software. The predicted locations of transmembrane domain (TMD) and signal peptide (SP) are indicated where applicable. N.D., not determined.

| Software | Prediction of protein type | Source |
| --- | --- | --- |
| TMHMM | Transmembrane protein (TMD: aa 10 - 29) | (4) |
| SMART | Transmembrane protein (TMD: aa 10 - 29) | (5) |
| SOSUI | Transmembrane protein (TMD: aa 11 - 33) | (6) |
| Phyre2 | Transmembrane protein (SP: aa 1 - 23; TMD: aa 46 - 61) | (7) |
| TMPred | Transmembrane protein (TMD: aa 13 - 29) | (8) |
| HMMTOP | Transmembrane protein (TMD: aa 4 - 25) | (9) |
| OCTOPUS | Transmembrane protein (TMD: aa 11 - 30) | (10) |
| Philius | Transmembrane protein (TMD: aa 8 - 29) | (11) |
| SCAMPI | Transmembrane protein (TMD: aa 9 - 29) | (12) |
| UniProt | Secreted protein (SP: aa 1 - 29) | (13) |
| DeepLoc | Secreted protein (SP: N.D.) | (14) |
| TOPCONS | Secreted protein (SP: aa 1 - 30) | (15) |
| PolyPhobius | Secreted protein (SP: aa 1 - 31) | (16) |
| SPOCTOPUS | Secreted protein (SP: aa 1 - 30) | (17) |
| TMSEG | Secreted protein (SP: aa 1 - 27) | (18) |

**Table S2.** Information about the antibodies used in this study.

| **Antibodies** | **Source** | **Identifier** |
| --- | --- | --- |
| Rabbit polyclonal anti-1700029I15Rik antibody | This study | N/A |
| Mouse monoclonal anti-1D4 antibody | A gift from Dr. Martin M. Matzuk | N/A |
| Mouse monoclonal anti-β-actin antibody | Abcam | ab6276 |
| Rabbit polyclonal anti-β-actin antibody | MBL International | PM053 |
| Rat monoclonal anti-ADAM1B | Laboratory of Masahito Ikawa | KS107-158 |
| Mouse monoclonal anti-ADAM3 antibody | Santa Cruz Biotechnology | sc-365288 |
| Mouse monoclonal anti-BASIGIN antibody | Santa Cruz Biotechnology | sc-46700 |
| Rabbit monoclonal anti-CANX antibody | Laboratory of Masahito Ikawa | 99051 |
| Rabbit monoclonal anti-CALR antibody | Laboratory of Masahito Ikawa | C1288 |
| Rabbit monoclonal anti-CALR3 antibody | Laboratory of Masahito Ikawa | B6530 |
| Rat monoclonal anti-CD46 antibody | Laboratory of Masahito Ikawa | KS107-116 |
| Rabbit monoclonal anti-CLGN antibody | Laboratory of Masahito Ikawa | 83170 |
| Rabbit monoclonal anti-COX4 antibody | Abcam | ab202554 |
| Mouse monoclonal anti-EQTN antibody | A gift from Dr. Kiyotaka Toshimori | N/A |
| Mouse monoclonal anti-FLAG antibody | Sigma-Aldrich | F1804 |
| Rabbit polyclonal anti-FLAG antibody | MBL International | PM020 |
| Rabbit polyclonal anti-GAPDH antibody | Cell Signaling Technology | 14C10 |
| Mouse polyclonal anti-GOLGIN97 antibody | Abcam | ab84340 |
| Rabbit polyclonal anti-HSP90B1 antibody | Cell Signaling Technology | 2104 |
| Rabbit polyclonal anti-HSPA5 antibody | Affinity BioReagents | PA1-014 |
| Rat monoclonal anti-IZUMO1 antibody | Laboratory of Masahito Ikawa | KS064-125 |
| Mouse monoclonal anti-KDEL antibody | StressGen | SPA-827 |
| Mouse monoclonal anti-P4HB antibody | Affinity BioReagents | MA3-019 |
| Rat monoclonal anti-PA antibody | FUJIFILM Wako | NZ-1 |
| Mouse monoclonal anti-PDIA3 antibody | Abcam | ab13506 |
| Rabbit polyclonal anti-SPACA1 antibody | Laboratory of Masahito Ikawa | B9167 |
| Rat monoclonal anti-SPACA4 antibody | Laboratory of Masahito Ikawa | KS139-281 |
| Rabbit polyclonal anti-SPACA6 antibody | This study | N/A |
| Rabbit polyclonal anti-SYCP1 antibody | Abcam | ab15090 |
| Chicken polyclonal anti-SYCP3 antibody | Laboratory of Hiroki Shibuya | N/A |
| Rabbit monoclonal anti-TOMM20 antibody | Abcam | ab186735 |
| Mouse monoclonal anti-Ubiquitin antibody | MBL International | MK-11-3 |
| Rabbit polyclonal anti-VDAC3 antibody | Proteintech | 14451-1-AP |
| Mouse monoclonal anti-ZP3R antibody | Invitrogen | MA1-10866 |
| Goat polyclonal anti-ZPBP1 antibody | A gift from Dr. Martin M. Matzuk | G176 |

**Table S3.**
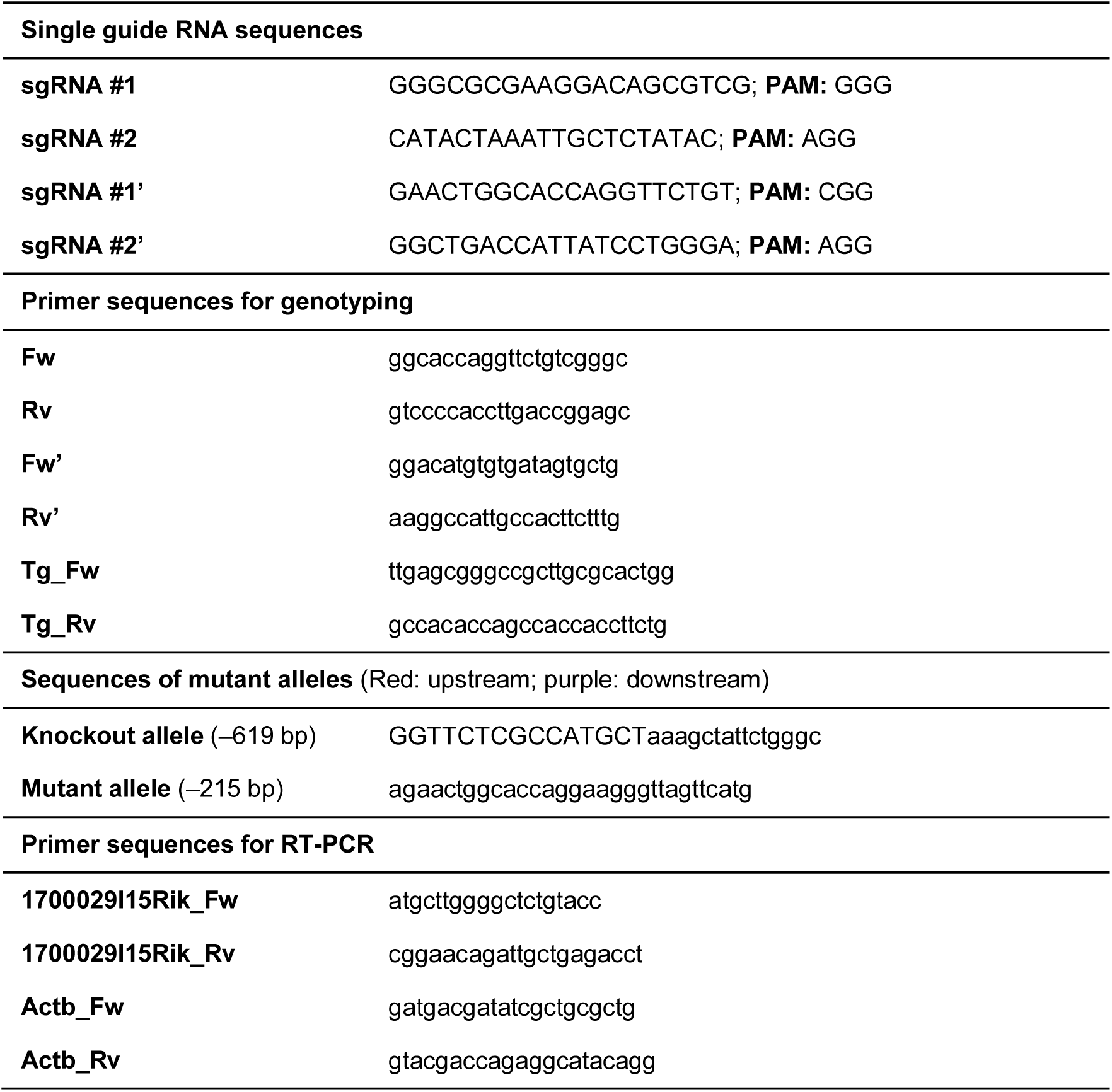
sgRNAs and primers used in this study.

### SI Figure Legends

**Fig. S1.**
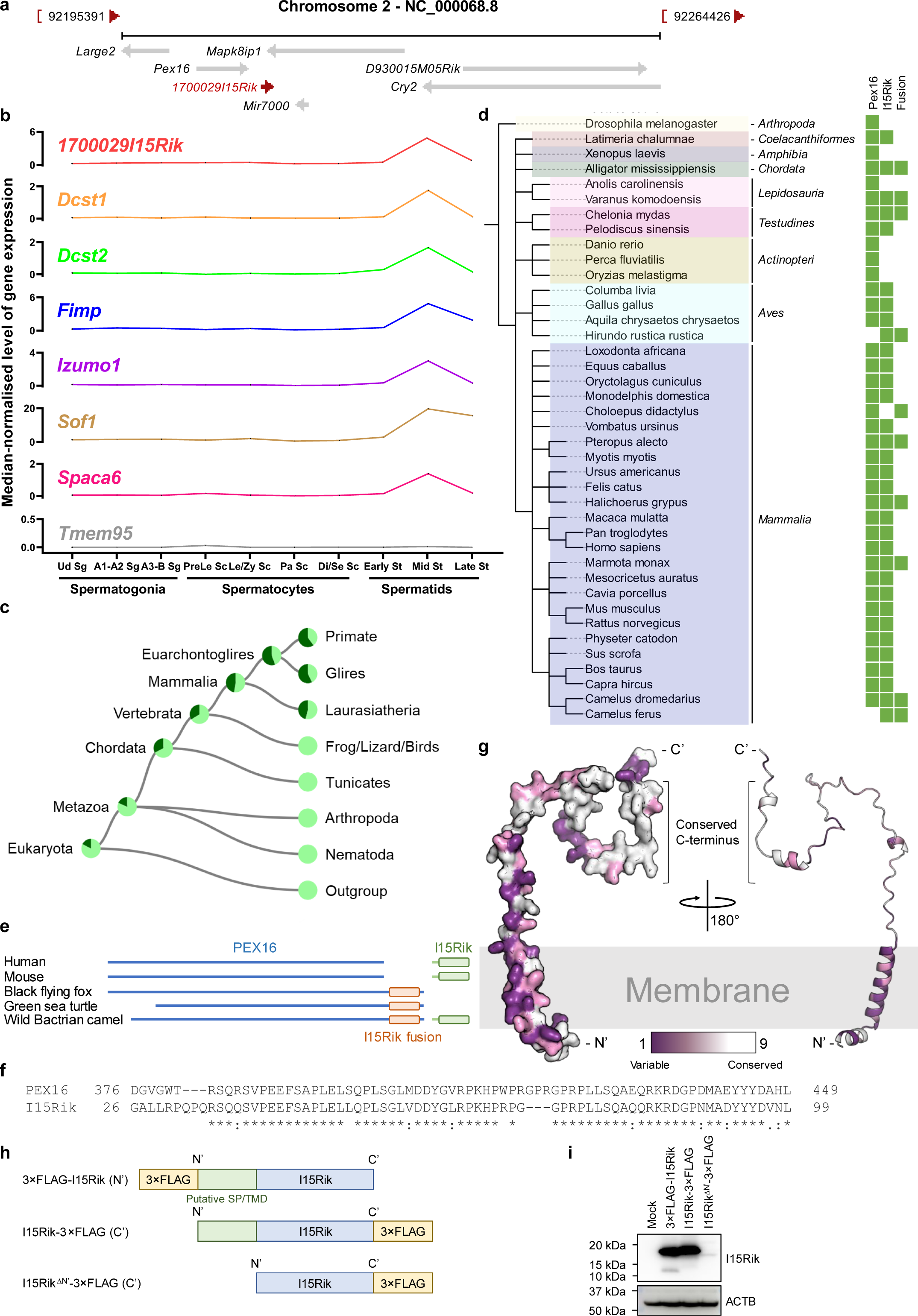
Characterisation of mouse *1700029I15Rik*. **a**, Chromosomal localisation of mouse *1700029I15Rik*. Adapted from the NCBI database. **b**, mRNA expression of *1700029I15Rik*, *Dcst1*, *Dcst2*, *Fimp*, *Izumo1*, *Sof1*, *Spaca6* and *Tmem95* in mouse spermatogenic cells based on the scRNA-seq datasets published previously (1). Ud Sg, undifferentiated spermatogonia; A1-A2 Sg, types A1 to A2 spermatogonia; A3-B Sg, types A3 to B spermatogonia; PreLe Sc, preleptotene spermatocytes; Le/Zy Sc, leptotene/zygotene spermatocytes; Pa Sc, pachytene spermatocytes; Di/Se Sc, diplotene/secondary spermatocytes; St, spermatids; Early St, early round spermatids; Mid St, mid-round spermatids; Late St, late round spermatids. **c**, Phylogenetic tree of *1700029I15Rik* generated by TreeFam (19, 20). **d**, Taxonomic tree depicting the presence and absence of 1700029I15Rik and PEX16 in various species. The tree was visualised by the iTOL software (21). **e**, Protein structures of PEX16 and 1700029I15Rik in various species. **f**, Alignment of the C-terminal sequences of woodchuck PEX16 and mouse 1700029I15Rik. **g**, Analyses of the evolutionary conservation of amino acid positions in 1700029I15Rik by the ConSurf software (22). **h**, Plasmids encoding epitope-tagged 1700029I15Rik for analysing the protein topology. 3×FLAG-I15Rik (N’), N-terminal 3×FLAG-tagged 1700029I15Rik; I15Rik-3×FLAG (C’), C-terminal 3×FLAG-tagged 1700029I15Rik; I15Rik^ΔN’^-3×FLAG (C’), N-terminus truncated, C-terminal 3×FLAG-tagged 1700029I15Rik. SP, signal peptide; TMD, transmembrane domain. **i**, Western blot analyses of recombinant expression of the epitope-tagged 1700029I15Rik in HEK293T cells. GAPDH was analysed in parallel as a loading control.

**Fig. S2.**
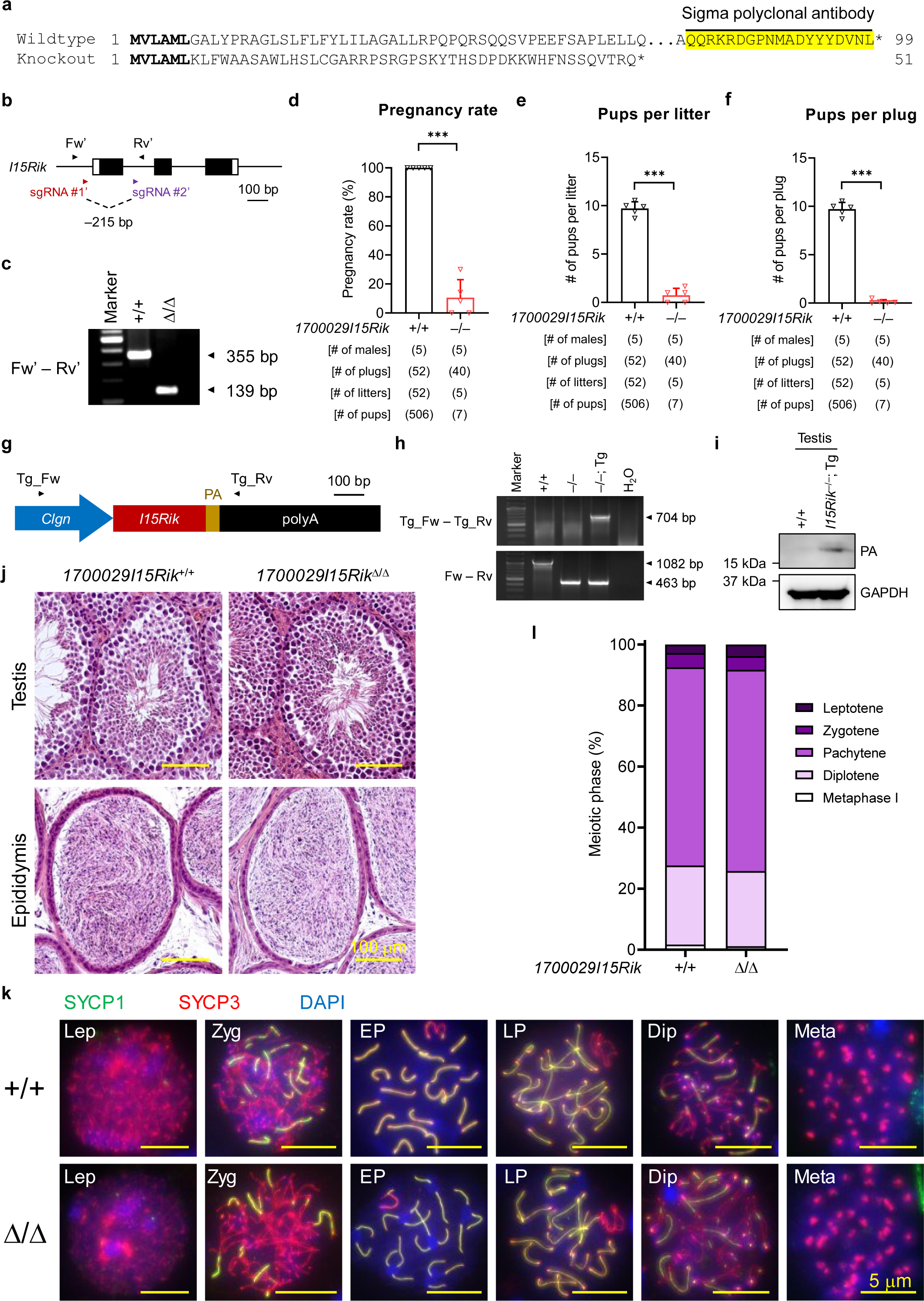
1700029I15Rik-deficient male mice are infertile but exhibit normal spermatogenesis. **a**, Amino acid sequences translated from the wildtype and knockout alleles. The sequences in bold are identical in the wildtype and knockout alleles. **b**, Schematics of generating the 1700029I15Rik mutant mouse line on the background of C57BL/6J by CRISPR/Cas9. **c**, Genomic PCR for distinguishing mice carrying the wildtype and mutant alleles using the forward (Fw’) and reverse (Rv’) primers flanking the first coding exon of *1700029I15Rik*. **d – f**, Analyses of the fecundity of wildtype and knockout males by fertility tests. The male fecundity is presented by pregnancy rate (the total number of litters over the total number of plugs), average number of pups per plug, or the average number of pups per litter. **g**, Schematics of transgenic expression using a linearised plasmid encoding a *Clgn* promoter, the ORF of *1700029I15Rik* (*I15Rik*), the PA tag, and a rabbit beta-globin polyadenylation (polyA) signal. **h**, Genomic PCR for identifying the PA-tagged *1700029I15Rik* transgene using the forward (Tg_Fw) and reverse (Tg_Rv) primers targeting the *Clgn* promoter and the polyA tail, respectively. **i**, Western blot analysis of transgenic expression of PA-tagged 1700029I15Rik in the testis. GAPDH was analysed in parallel as a loading control. **j**, Histological analyses of wildtype and mutant testes and epididymides. **k**, Immunostaining of wildtype and mutant spermatocyte chromosome spreads. The transverse filament and axial element of meiotic chromosomes were visualised by probing with anti-SYCP1 (green) and SYCP3 (red), respectively. The chromatin was stained with DAPI in blue.

**Fig. S3.**
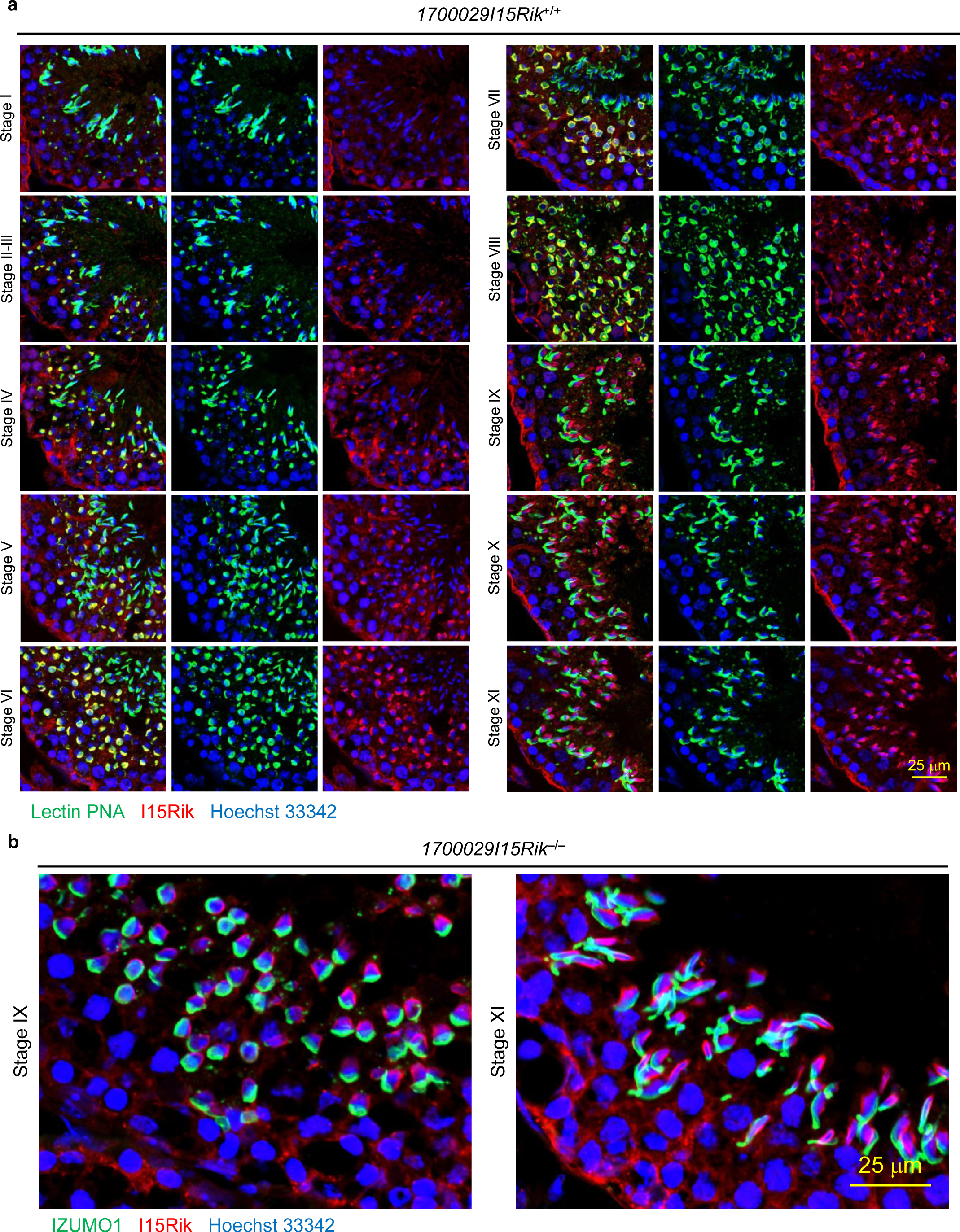
1700029I15Rik is localised to the acrosome in early round spermatids and absent in the elongating spermatids. **a**, Detailed analysis of the localisation of 1700029I15Rik (red) in each spermatogenic stage by immunohistochemistry. The acrosome was stained by lectin PNA in green and the cell nuclei was visualised by Hoechst 33342 in blue. **b**, Non-specific recognition of manchette by the 1700029I15Rik antibody as revealed by immunostaining of the knockout testis cryosections.

**Fig. S4.**
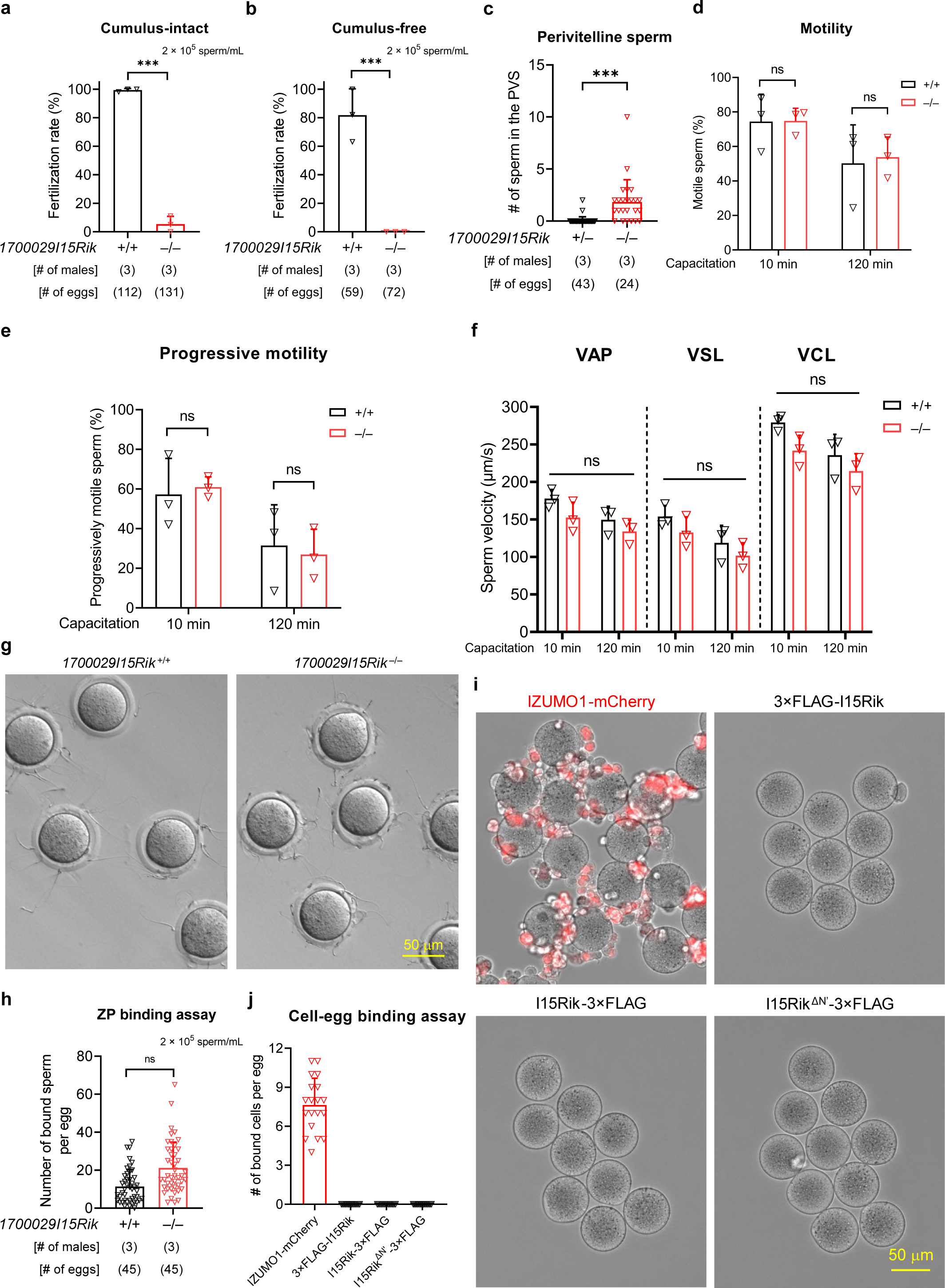
*1700029I15Rik* knockout sperm show impaired fertilising ability but normal motility and ZP binding ability. **a – b**, IVF using cumulus-intact and cumulus-free wildtype eggs. **c**, Analysis of the numbers of perivitelline sperm in the eggs retrieved from superovulated female mice that had been paired with wildtype or knockout males. **d – f**, Analyses of sperm motility, progressive motility, and velocity at 10 and 120 min of incubation in the TYH medium by CEROS II. VAP, average path velocity; VSL, straight line velocity; VCL, curvilinear velocity. **g – h**, Analysis of the ZP binding ability of wildtype and knockout spermatozoa. **i - j**, Analyses of the oolemma binding ability of HEK293T cells transiently expressing mCherry-tagged IZUMO1 (red) and 1700029I15Rik.

**Fig. S5.**
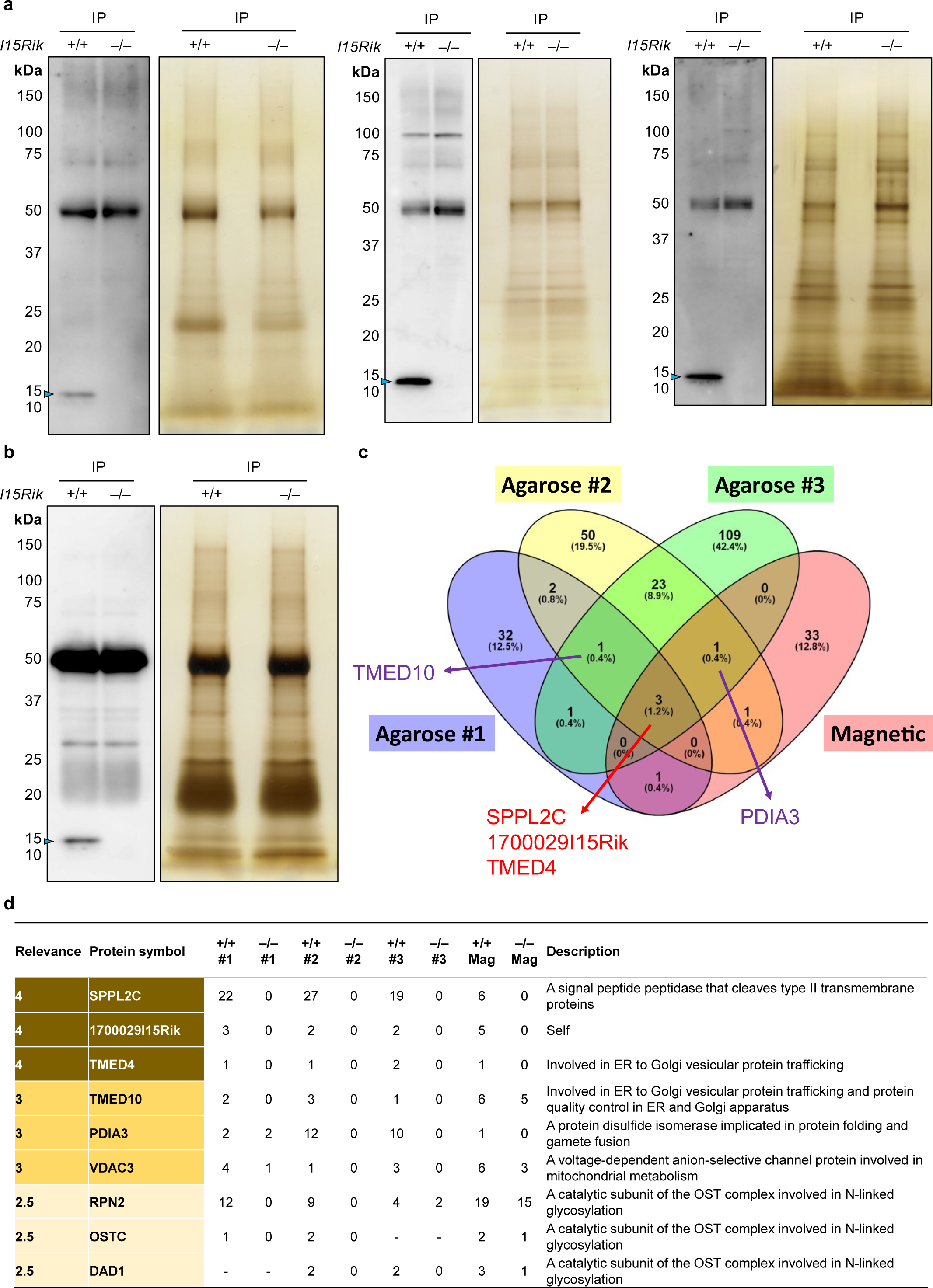
1700029I15Rik interacts with proteins implicated in N-linked glycosylation, disulfide isomerisation, and ER–Golgi vesicular trafficking. **a**, Western blot and silver staining analyses of proteins immunoprecipitated using the polyclonal 1700029I15Rik antibody covalently crosslinked to the agarose resin. Blue arrowheads indicate 1700029I15Rik protein band is immunodetected in the wildtype but absent in the knockout sample. **b**, Western blot and silver staining analysis of proteins immunoprecipitated using 1700029I15Rik antibody-conjugated magnetic beads. The antibody was not covalently crosslinked to the magnetic beads. Blue arrowhead indicates 1700029I15Rik protein band is immunodetected in the wildtype but absent in the knockout sample. **c**, Venn diagram of proteins concurrently identified in the four trials of the co-IP/MS experiments. Only the proteins specifically detected in the wildtype and absent in the knockout sample were analysed. **d**, Normalised total spectra of proteins detected in each co-IP/MS experiment. The specificity of the protein interactions was scored based on the numbers of detected spectra [1 for specific detection in the wildtype sample; 0.5 for fold changes (wildtype over knockout) ≥ 2; 0 for fold changes < 2].

**Fig. S6.**
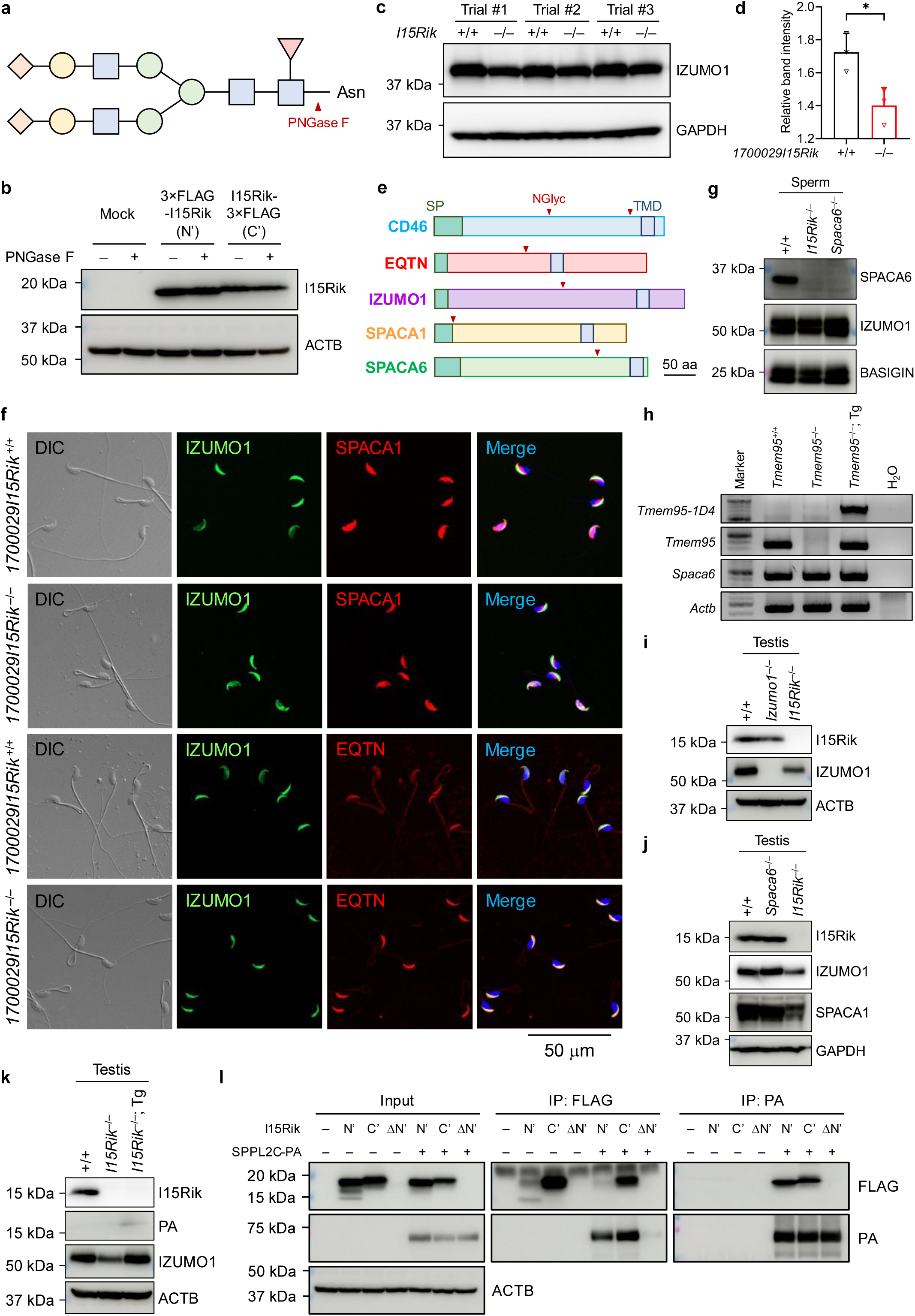
Ablation of 1700029I15Rik results in downregulation of multiple acrosomal membrane proteins in the knockout testis. **a**, Removal of the asparagine (Asn)-linked glycan sidechain by PNGase F. Blue squares, green circles, yellow circles, and orange diamonds indicate N-acetylglucosamine, galactose, mannose, and sialic acid, respectively. **b**, N-glycosylation analysis of 1700029I15Rik transiently expressed in HEK293T cells. ACTB was analysed as a loading control. **C – d**, Western blot analyses of the amounts of IZUMO1 in the wildtype and knockout testes. GAPDH was analysed in parallel as a loading control. The intensities of IZUMO1 and GAPDH protein bands were measure by ImageJ and the band intensities of IZUMO1 relative to GAPDH were calculated. **e**, Protein structures of CD46, EQTN, IZUMO1, SPACA1, and SPACA6. The signal peptide (SP), transmembrane domain (TMD), and N-glycosylation (NGlyc) sites are highlighted. **f**, Immunocytochemistry analyses of IZUMO1, SPACA1, and EQTN in wildtype and knockout spermatozoa. **g**, Western blot analysis of SPACA6 in wildtype, *1700029I15Rik* knockout, and *Spaca6* knockout spermatozoa. IZUMO1 and BASIGIN were analysed in parallel. **h**, RT-PCR analyses of the mRNA expression of the *Tmem95-1D4* transgene, *Tmem95*, and *Spaca6* in wildtype, *Tmem95*^−/−^, *Tmem95*^−/−^; Tg testes. The mRNA expression of *Actb* was analysed in parallel as a loading control. **i – j**, Western blot analyses of 1700029I15Rik in *Izumo1*^−/−^ and *Spaca6*^−/−^ testes. IZUMO1 and SPACA1 were analysed in parallel. ACTB or GAPDH was analysed as a loading control. **k**, Western blot analyses of 1700029I15Rik, 1700029I15Rik-PA, and IZUMO1 in wildtype, *1700029I15Rik*^−/−^, and *1700029I15Rik*^−/−^; Tg testes. ACTB was analysed as a loading control. **l**, Co-IP analyses of interactions between SPPL2C and 3×FLAG-tagged 1700029I15Rik in HEK293T cells. N’, 3×FLAG-1700029I15Rik; C’, 1700029I15Rik-3×FLAG; ΔN’, 1700029I15Rik^ΔN’^-3×FLAG. Protein complexes were immunoprecipitated by FLAG or PA antibody. ACTB were analysed as a loading control for the input samples.

**Fig. S7.**
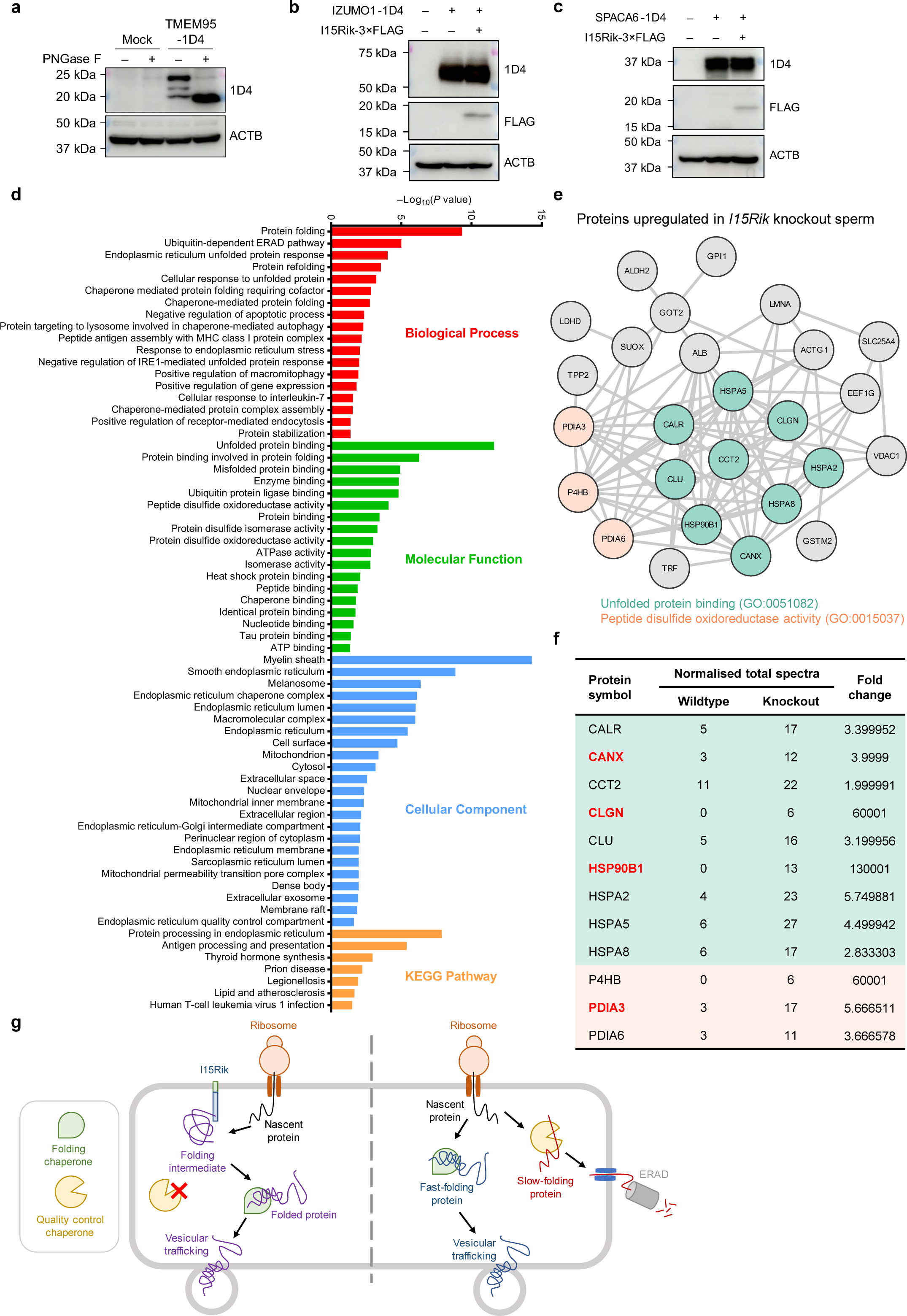
Depletion of 1700029I15Rik leads to upregulation of ER chaperones involved in the ERAD pathway in the sperm. **a**, N-glycosylation analysis of TMEM95 transiently expressed in HEK293T cells. ACTB was analysed as a loading control. TMEM95 has two N-linked glycosylation sites. From top to bottom, the three protein bands in the untreated sample represent fully glycosylated, partially glycosylated, and non-glycosylated TMEM95. **b**, Co-expression of 1700029I15Rik and IZUMO1 in HEK293T cells. **c,** Co-expression of 1700029I15Rik and SPACA6 in HEK293T cells. **d - e**, GO, KEGG (23), and STRING (24, 25) analyses of proteins upregulated in the knockout spermatozoa. **f**, Normalised total spectra of the ER chaperones upregulated in the knockout spermatozoa. The proteins highlighted in red were upregulated in the knockout sperm as confirmed by the western blot analyses (Fig. 5f). **g**, A diagram illustrating a hypothetical model of 1700029I15Rik-mediated protein processing. Nascent proteins are modified by 1700029I15Rik into folding intermediates so as to be exempted from being destructed the ERAD pathway (left). In the absence of 1700029I15Rik, slow-folding proteins undergo degradation via ERAD, whereas fast-folding molecules are escaped from the targeted destruction (right).

